# ChemIntelligence Enables Antibody-Free, Ultra-Low-Input Profiling of Lysine Lactylation and Diverse Acyl-Proteomes

**DOI:** 10.64898/2026.08.28.746934

**Authors:** Chang Shao, Zimeng He, Qi Yuan, Victor-George Giurcoiu, Xiangyu He, Xu Cao, Haoran Huang, Yuebo Zhang, Yueyang Zhang, Dexiang Wang, Qihe Jiang, Zizhang Guo, Haiping Hao, Mathias Wilhelm, Hui Ye

## Abstract

Lysine acylations, including lactylation (Klac), are pivotal regulators of cellular physiology. However, their analysis is currently bottlenecked by antibody enrichment strategies that suffer from sequence bias and require milligram-scale protein inputs, severely precluding the profiling of scarce clinical biopsies and rare cell populations. Here we present **ChemIntelligence**, an acyl-NHS chemistry-empowered derivatization strategy that rapidly generates unprecedented acylation-specific spectral libraries, exemplified by over 2.5×10^5^ human Klac peptides, enabling cross-species reference atlases. Integrated with Prosit-based rescoring, these libraries substantially increase Klac identifications across diverse proteomic datasets. Leveraging this spectral resource, we devised **ChemIntelligence Scope**, a reproducible, multiplexed parallel reaction monitoring (PRM) platform that quantifies hundreds of Klac peptides per injection from as little as ∼200 ng of cell lysates, clinical biopsies, and even true single cells—revealing functional Klac signatures inaccessible to conventional methods. The ChemIntelligence pipeline also extends seamlessly to lysine nicotinylation, underscoring its broad adaptability for discovering and profiling new acylations. Together, these chemical and computational advances establish a scalable, antibody-free framework for acyl-proteome mapping that overcomes input constraints and enables deep functional insights from otherwise intractable biological samples.

## INTRODUCTION

The ε-amino group of lysine is intrinsically reactive, making it a privileged target for a wide spectrum of acylation events. Over the past decade, advances in proteomics have enabled reliable detection of numerous lysine acylations—including crotonylation^1^, succinylation^2^, malonylation^3^, glutarylation^4^, 2-hydroxyisobutyrylation^5^, β-hydroxybutyrylation^6^, benzoylation^7^, propionylation^8^, and Klac^9^. These discoveries have transformed lysine acylation into one of the fastest-expanding PTM families. Among these, Klac has attracted particular attention for its emerging roles in transcriptional regulation, DNA repair, immune responses, and tumor biology^10,11^.

Despite this progress, lysine acylations including Klac remain predominantly investigated via antibody-based enrichment^1–7,9^. This approach faces major upstream barriers: scarce antibody availability, high antibody production costs, long development timelines^12^, and intrinsic sequence biases that capture only a fraction of the modification landscape^13^. Generating antibodies for chemically subtle or rare acyl groups is particularly challenging, impeding systematic discovery of newly identified or hypothesized PTMs. Even when antibodies exist, downstream workflows require mg-scale protein inputs to achieve detectable yields^1–7,9,12^, making them impractical for low-input biosamples including rare primary cell subsets, clinical biopsies or single-cell isolates. Together, these constraints limit sensitivity, reproducibility, and—critically—the ability to uncover acylations from scarce or heterogeneous specimens.

Targeted proteomic approaches—spanning from early selected reaction monitoring (SRM) and multiple reaction monitoring (MRM) to the more recent PRM—offer highly specific and sensitive quantification of peptides from minimal samples, making them particularly attractive for profiling precious clinical specimens^14–19^. The advent of synthetic peptide libraries, and their associated spectral datasets has transformed targeted assay development, enabling serial SRM/MRM–based, large-scale proteome profilin^20–24^. Modern PRM platforms, such as the Stellar quadrupole–linear ion trap (Q-LIT) mass spectrometer, further increase analytical throughput by combining high multiplexing capacity with reproducible, large-scale detection in streamlined assays^17–19^. However, this synthetic-library enabled targeted proteomics paradigm has rarely been extended to lysine acylations, primarily due to the lack of modification-specific peptide reference libraries. Even lysine acetylation (Kac)—the most extensively studied acylation—is represented by merely ∼200 reference peptides in the ProteomeTools resource^21,25^, a scale far too limited to support comprehensive PRM assay development. This scarcity is even more pronounced for other acylations, particularly newly characterized ones such as Klac, for which no libraries are available. The field therefore remains constrained by antibody-dependent methods, hindering the exploration of acyl-proteomes in clinically relevant settings.

Here we address this gap by developing **ChemIntelligence**, a generalizable chemical derivatization strategy that employs acyl-NHS ester chemistry to rapidly generate the largest, acylation-specific spectral libraries across human and other species. This approach converts complex proteomes into extensive acyl-peptide repertoires within days at minimal cost (∼10 USD per 500 mg reagent), bypassing labor-intensive peptide synthesis and producing datasets suitable for multiple downstream applications such as deep learning model training and targeted PRM assay development (**Fig. 1a**, left panel).

**Figure 1.**
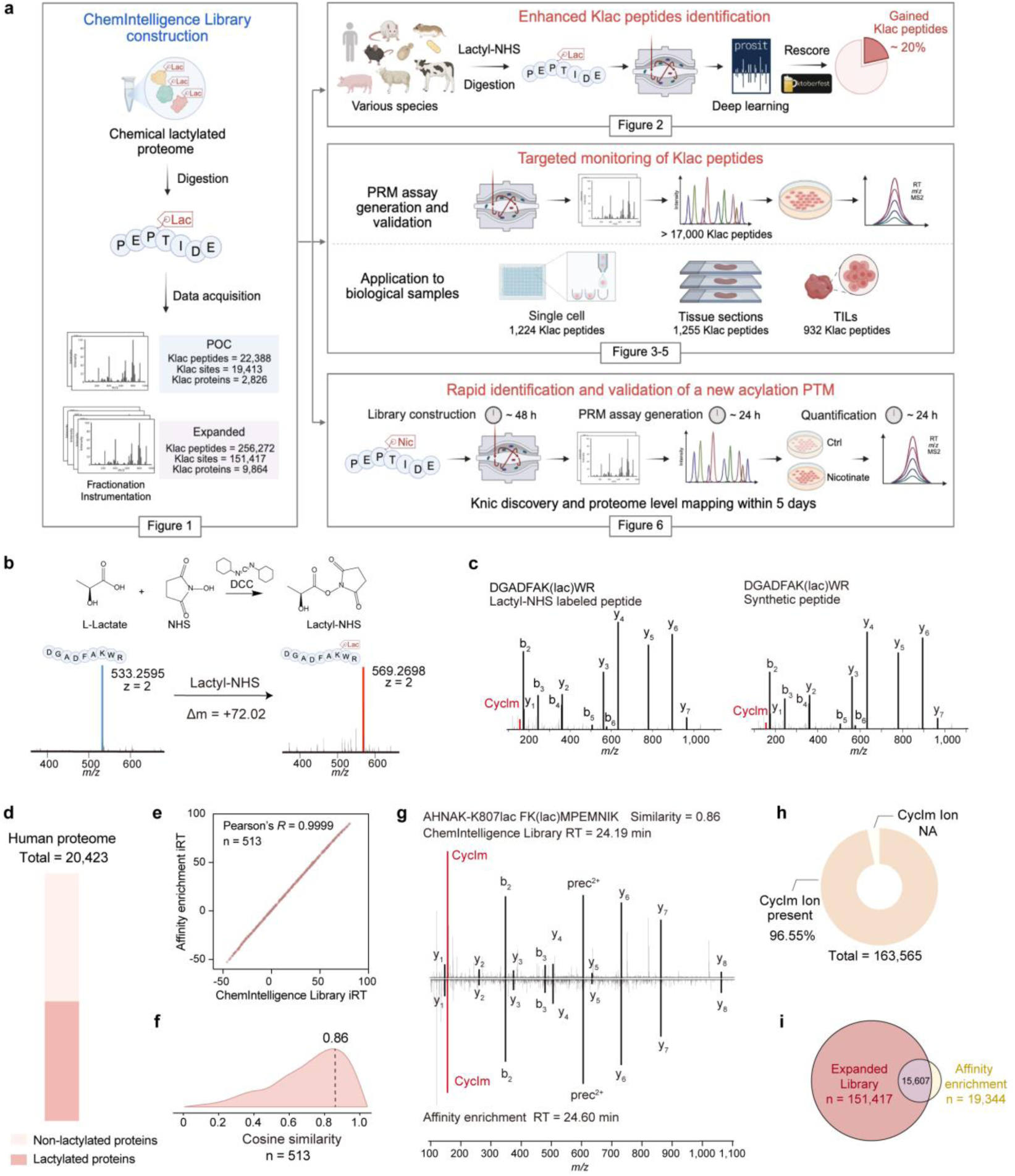
Large-scale Klac peptide library constructed via the ChemIntelligence strategy enables antibody-free, scalable lactylproteome mapping in low-input samples. (a) Overview of the development of the ChemIntelligence Library and its diverse functions in acyl-proteome mapping using Klac as an exemplary. The left panel depicts its design and construction; the right panels illustrate downstream applications, including rescoring enhanced Klac identification (Fig. 2), PRM-based ChemIntelligence Scope analysis of bulk, low-input, and clinical samples (Figs. 3–5), and extension to novel modifications such as nicotinylation (Fig. 6). (b) Top, synthetic scheme for generating lactyl-NHS ester. NHS, *N*-hydroxysuccinimide; DCC, *N*, *N*’-dicyclohexylcarbodiimide. Bottom, MS spectra of a model peptide from ALDOA before (left) and after (right) chemical lactylation introduced by lactyl-NHS. (c) MS/MS spectra comparison of the chemically generated Klac peptide (left) and the corresponding synthetic standard (right). (d) Broad proteome coverage of Klac in the ChemIntelligence Library, mapped across the human proteome. (e) Correlation of iRTs for Klac peptides detected in the ChemIntelligence Library and those from the affinity-enriched HeLa cells treated with lactate (Pearson’s *R* > 0.99), confirming retention time fidelity. (f) Distribution of cosine similarity comparing MS/MS spectra of matched Klac peptides in (**e**). Lower similarity scores (< 0.86) are attributed to reduced spectral qualify for enriched endogenous Klac peptide (related to **Supplementary Fig. S5b**). (g) Mirror plot comparison of the representative peptide FK(lac)MPEMNIK derived from the ChemIntelligence Library (top) and affinity-enriched HeLa cells (bottom), corresponding to the apex in (**f**). (h) Frequency of CycIm ion detection in ChemIntelligence Library of Klac peptides acquired using Orbitrap instruments. (i) Venn diagram comparing Klac peptide coverage in the ChemIntelligence Library with published affinity-enriched lactylproteome datasets, highlighting identification of previously unreported Klac sites.

In this study, we demonstrate the platform’s utility for Klac by constructing the **ChemIntelligence Library**, comprising over 256,000 Klac peptides for the human proteome and over 54,000 across other taxa. Leveraging this comprehensive reference, we implemented a deep learning-based Prosit–Oktoberfest rescoring workflow^26,27^ that markedly improves Klac peptide identification across diverse proteomic datasets, including re-analysis of existing antibody-enrichment experiments (**Fig. 1a**, upper right panel). Building on this spectral resource, we further developed the **ChemIntelligence Scope** strategy for Klac, a PRM-based platform that enables reproducible, scalable, antibody-free Klac detection and quantification from as little as ∼200 ng of HeLa digest, with sensitivity sufficient to identify hundreds of Klac peptides in single-cell isolates within a 30-minute gradient. When applied to clinical tumor biopsies and tumor-infiltrating T cells, this approach uncovered functional Klac signatures in clinically relevant contexts that are inaccessible to antibody-based methods (**Fig. 1a**, middle right panel). To highlight the platform’s generality, we further adapted the ChemIntelligence pipeline to lysine nicotinylation^28^ (Knic), enabling rapid spectral-library creation and proteome-wide mapping of this recently described acylation within five days (**Fig. 1a**, lower right panel).

Together, the combination of ChemIntelligence library construction, deep learning-based rescoring, and targeted PRM workflow transforms lysine acyl-proteome analysis from an antibody-dependent paradigm to a standardized and scalable framework. This shift enables the functional exploration of lysine acyl-proteome in previously inaccessible yet biologically critical settings, from single cells to limited clinical biopsies, positioning our approach as a powerful enabler for deciphering their functional biology.

## RESULTS

### ChemIntelligence empowers rapid, large-scale construction of Klac spectral libraries

To build the ChemIntelligence Library, using Klac as an exemplary acylation, we designed and synthesized lactyl-NHS (**Fig. 1b, Supplementary Fig. S1**)—a core Klac modifier inspired by the structural concept of the acetylation inducer acetyl-NHS^29^. We first demonstrated that lactyl-NHS can efficiently introduce lactylation onto free lysines in peptides (**Fig. 1b-c**) and, following concentration optimization, onto lysines in model proteins (**Supplementary Fig. S2a-b**). Digestion of these chemically lactylated proteins yielded a diverse repertoire of Klac peptides (**Supplementary Fig. S2c-f**), eliminating the need for traditional peptide synthesis. We next benchmarked the lactylation performance of lactyl-NHS for complex biological samples (**Supplementary Fig. S3**) and used it to construct a Klac peptide reference library via the ChemIntelligence strategy. Chemical lactylation of HeLa cell lysates followed by deep proteomic analysis on the Orbitrap Astral analyzer resulted in a proof-of-concept (POC) ChemIntelligence Library comprising 22,388 Klac peptides and 19,413 lysines (**Fig. 1a, Supplementary Table 1**). This resource was further expanded by incorporating fractionation and complementary MS platforms, resulting in a compendium of 256,272 Klac peptides mapped to 9,864 proteins, with 227,062 identified as unique sequences (**Fig. 1a** and **Supplementary Table 2**). The resulting library offers broad, unbiased coverage with high efficiency and low cost (**Fig. 1d, Supplementary Fig. S4a**). Physicochemical profiling confirmed that the chemically-generated Klac peptides exhibit no bias in hydrophilicity, molecular weight, protein class or secondary structure compared to the human proteome (**Supplementary Fig. S4b-e**).

Next, we evaluated the fidelity of Klac peptides in the chemically-generated ChemIntelligence Library by benchmarking them against endogenous counterparts. Indexed retention time (iRT) values showed strong concordance with those from affinity-enriched Klac peptides in HeLa cells (Pearson’s *R* > 0.99), confirming consistent chromatographic behavior (**Fig. 1e**). MS/MS spectra further demonstrated high similarity, with a peak cosine similarity of ∼0.86 (**Fig. 1f-g**, **Supplementary Fig. S5**). Additionally, 96.55% of ChemIntelligence peptides produced the diagnostic cyclic immonium (CycIm) ion^30^ characteristic of Klac (**Fig. 1h, Supplementary Table 2**), reinforcing the spectral integrity of the library.

With validated fidelity, the ChemIntelligence Library offers a cost-effective, scalable alternative to synthetic peptide resources to support large-scale, targeted proteomics workflows, capturing 89.7% previously unreported Klac sites (**Fig. 1i, Supplementary Table 3**). Compared to reported affinity-enriched lactylproteome, the library shows no bias in sequence or hydrophilicity (**Supplementary Fig. S6a-b**) and enhances coverage of cytosolic and mitochondrial proteins through Gene Ontology (GO) cellular component (CC) analysis (**Supplementary Fig. S6c**). Chemically generated for broad, unbiased representation, our library is publicly available and provides a valuable resource for the proteomics community. Its unique advantages over traditional synthetic peptide libraries^20,21,23^ also includes greatly reduced cost, simplified workflow and the ease to quickly adapt to other lysine acylations.

### Prosit-Lac–Oktoberfest rescoring unlocks deeper identification of Klac peptides

Previous modification-specific peptide reference libraries—such as those for phosphorylation^23,24^ or Kac^21,25^—have been generated synthetically to support computational proteomics workflows. To fully leverage the available data and enable extrapolation to other organisms, we aimed to train predictive models for Klac peptides. Given that Klac peptides alone might be insufficient, we complemented the spectral data from the Klac ChemIntelligence Library, acquired on Orbitrap Eclipse Tribrid and Orbitrap Astral instruments, with the PROSPECT^31^ dataset—post-annotated ProteomeTools^21^ spectra acquired on Orbitrap Fusion Lumos Tribrid originally used to train Prosit^26^. Using this combined, diverse dataset, two Prosit-Lac prediction models were trained: one for fragment-ion intensities and one for iRT (**Supplementary Fig. S7; Methods**). The intensity model accurately reproduced spectra for both unmodified and Klac-modified peptides, with median spectral angles of 0.90 and 0.84, respectively (**Fig. 2a**). Similarly, the retention time model also showed high accuracy (Pearson’s *R* = 0.98; 95% ΔiRT = 12.09; **Fig. 2b**).

**Figure 2.**
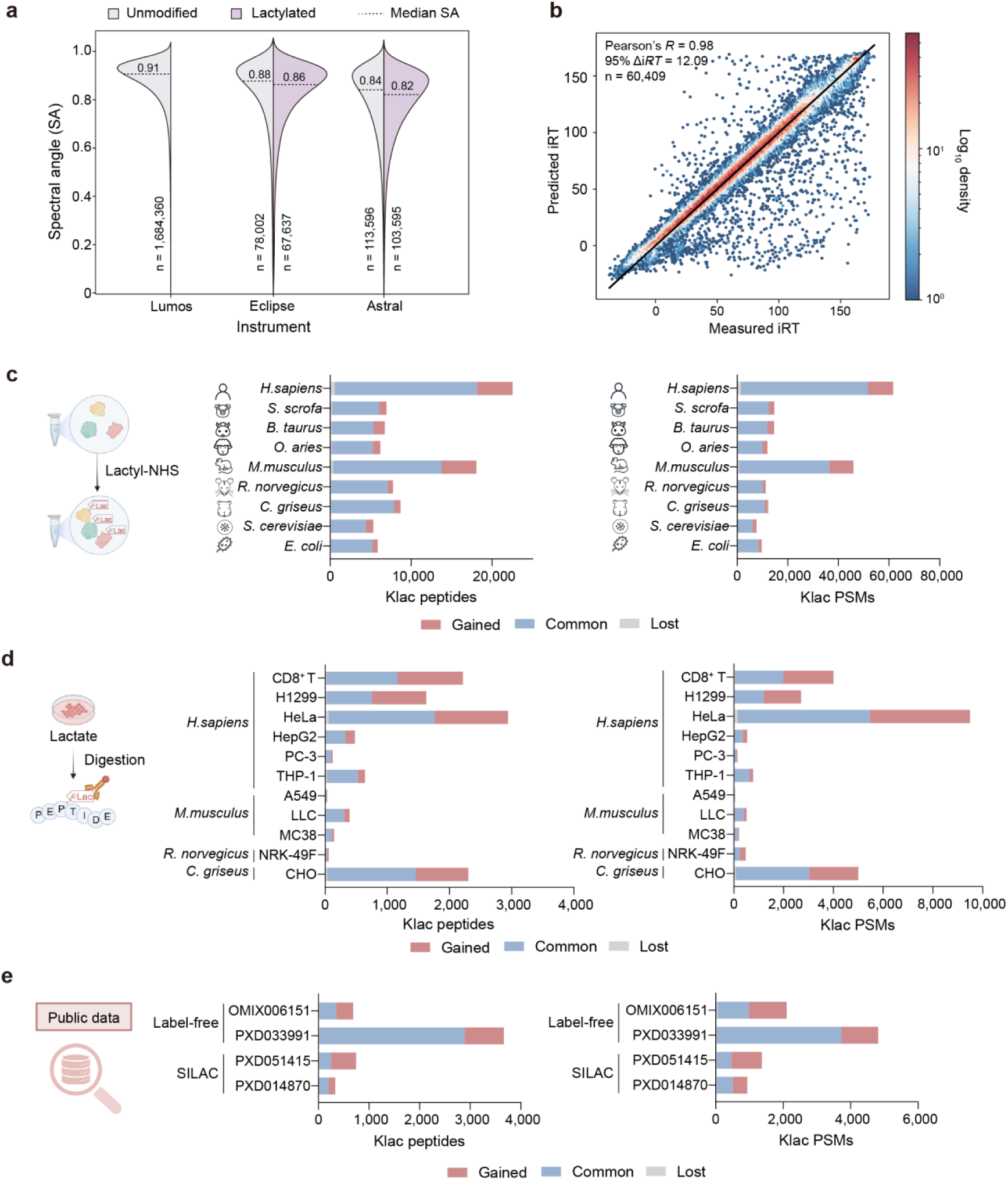
ChemIntelligence Library–assisted Prosit–Oktoberfest rescoring workflow improves identification of Klac peptides. (a) Violin plots depicting spectral angle distributions between Prosit-Lac–predicted and experimentally measured spectra in the test set. Data from unmodified peptides on the Orbitrap Lumos (left) are from the PROSPECT dataset, while data from both unmodified (grey) and lactylated (purple) peptides on the Eclipse (middle) and Astral (right) instruments were from expanded Klac ChemIntelligence Library. Dashed lines indicate medians. (b) Scatter density plot of the correlation between Prosit-Lac–predicted and experimentally measured iRTs for the test set. The color scale indicates point density on a Log_10_ scale. (c) Bar charts illustrating the numbers of gained (red), common (blue), and lost (grey) Klac peptides and PSMs after Oktoberfest rescoring of cross-species ChemIntelligence Libraries constructed for Klac, benchmarked against the original search results. (d) Bar charts illustrating the numbers of gained (red), common (blue), and lost (grey) Klac peptides and PSMs after Oktoberfest rescoring of affinity-enriched samples from lactate-treated cell lines of human and animal species, benchmarked against the original search results. (e) Bar charts illustrating the numbers of gained (red), common (blue), and lost (grey) Klac peptides and PSMs after Oktoberfest rescoring of publicly available affinity-enriched lactylproteome datasets, benchmarked against the original search results.

The trained models are publicly available on Koina^32^ and were integrated into our previously established Oktoberfest rescoring workflow^27^ to re-evaluate peptide-spectrum matches (PSMs), classifying Klac identifications as gained, common, or lost relative to the original database search results. Applied to the human Klac library generated from HeLa cell lysates, Prosit-Lac rescoring markedly improved Klac identification rates, increasing peptide and PSM counts by 24% and 19%, respectively (**Fig. 2c, Supplementary Fig. S8a**). To assess cross-species performance, Klac ChemIntelligence Libraries were built for 8 additional taxa beyond human, spanning mammals (*Sus scrofa*, *Bos taurus*, *Ovis aries*, *Mus musculus*, *Rattus norvegicus* and *Cricetulus griseus*) and microbes (*Saccharomyces cerevisiae* and *Escherichia coli*). Across these unenriched, Klac-rich datasets, the rescoring workflow delivered mean increases of 18% for peptides and 18% for PSMs (**Fig. 2c, Supplementary Fig. S8a**).

In affinity-enriched, endogenous Klac datasets acquired from human and animal cell lines, improvements were even greater, averaging 61% for peptides and 66% for PSMs (**Fig. 2d**, **Supplementary Fig. S8b**). Application to publicly available affinity-enriched datasets from varied Orbitrap platforms further confirmed its broad applicability: label-free datasets (OMIX006151, PXD033991) and SILAC datasets (PXD051415, PXD014870) showed mean increases of 94% (peptides) and 105% (PSMs) (**Fig. 2e**, **Supplementary Fig. S8c**).

In summary, integration of the Klac-specific Prosit-Lac predictors with Oktoberfest rescoring framework consistently expanded Klac peptide coverage across species, sample types, and acquisition strategies. This workflow supports both *de novo* Klac discovery and retrospective mining in diverse proteomics datasets, and can be readily applied to enhance identification rates for other acyl-proteomes using acyl-specific ChemIntelligence Libraries. All required tools are open source and provide pre-built pipelines for model training, prediction, and rescoring.

### ChemIntelligence Scope enables antibody-free mapping of lactylproteome in low-input biosamples

Conventional antibody-based Klac enrichment requires mg-scale protein input and prolonged workflows (**Fig. 3a**), limiting its applicability to scarce biological samples. To overcome these constraints, we developed the ChemIntelligence Scope strategy for Klac profiling—an antibody-free, high-throughput, targeted PRM workflow, powered by our Klac ChemIntelligence Library (**Fig. 3b**, upper panel). Implemented on a Q-LIT platform with serial PRM assays, ChemIntelligence Scope bypasses antibody dependence and enables direct interrogation of Klac from low-input and even single-cell samples, including HeLa cells, tumor biopsies, and infiltrating T cells (**Fig. 3b**, lower panel).

**Figure 3.**
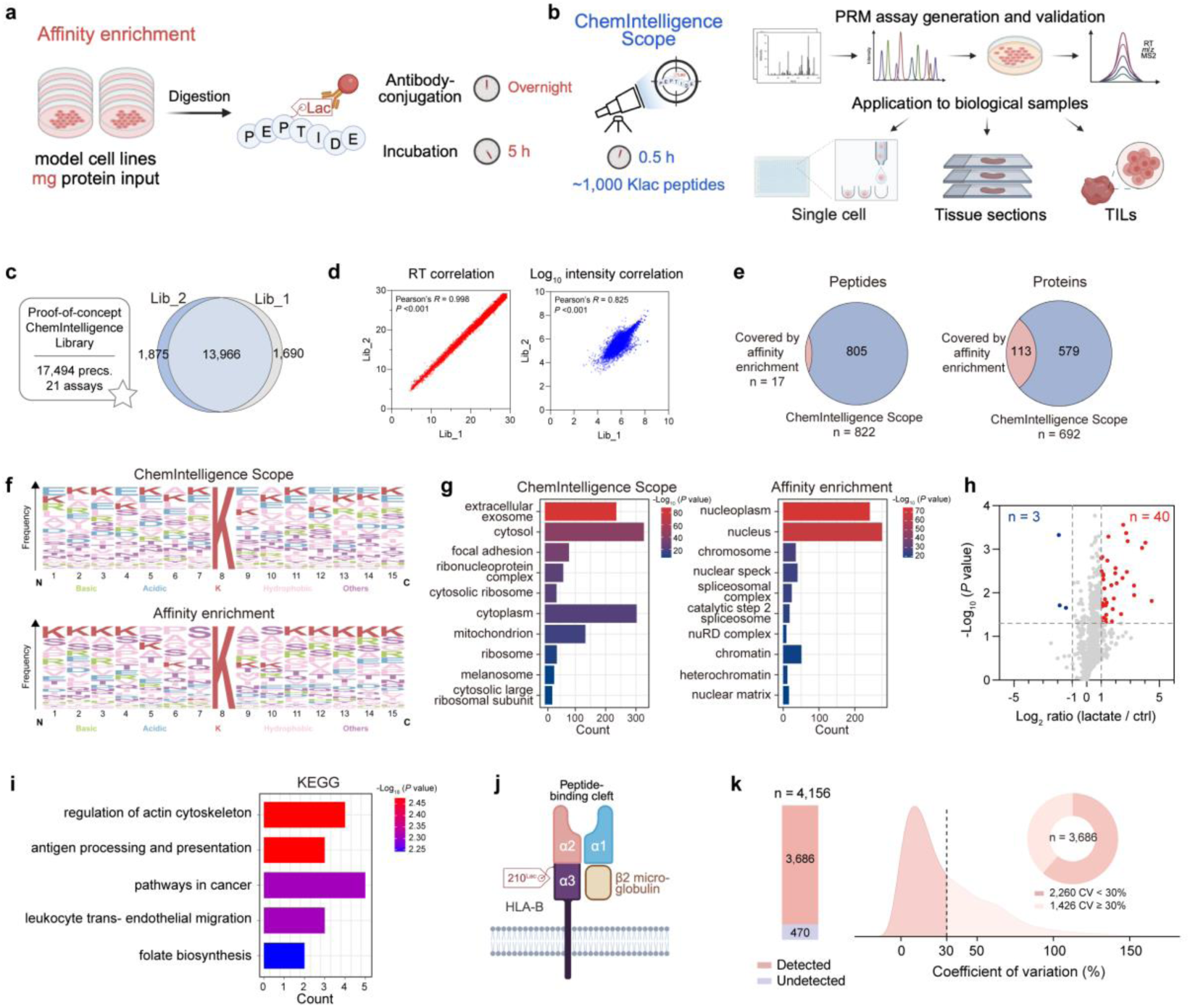
ChemIntelligence Scope assays were built based on ChemIntelligence Library, and enable targeted monitoring of large-scale Klac peptides from biosamples. (a) Workflow of the classic affinity enrichment strategy, involving overnight immune-conjugation followed by enrichment of Klac peptides from mg-scale protein inputs. (b) Workflow of the PRM-based ChemIntelligence Scope strategy, starting with ChemIntelligence spectral library construction, followed by targeted PRM assay development for sensitive, enrichment-free quantification from low-input samples, including single HeLa cells, tumor biopsy sections, and primary T cells. (c) Overlap of Klac peptides detected using the same PRM assay via the ChemIntelligence Scope approach in two experiments conducted five months apart. Following exclusion of peptides with truncated XICs from the analysis in Skyline, detected peptides were highly reproducible across the two batches. (d) High correlations of retention times (Pearson’s *R* > 0.99) and intensities (Pearson’s *R* > 0.82) were observed for shared Klac peptides detected across the two batches shown in panel (**d**). (e) Klac peptides and proteins quantified by ChemIntelligence Scope showed partial overlap with those detectable by affinity enrichment. (f) Sequence logos generated using WebLogo showing amino acid frequency surrounding lactylated sites in Klac peptides quantified by ChemIntelligence Scope (top) and by affinity enrichment (bottom). (g) Bar charts of GO enrichment analysis showing cellular component (CC) terms of lactylated proteins quantified by ChemIntelligence Scope (left) and by affinity enrichment (right). (h) Volcano plot of Klac peptides quantified by ChemIntelligence Scope, illustrating lactate treatment-associated changes in HeLa cells. Significantly altered peptides were defined as lactate/control ratio > 2 or < 0.5, *P* value < 0.05 by ratio paired *t*-test (n=3 biological replicates/group) and are highlighted in red (upregulated) and blue (downregulated). (i) Bar chart of KEGG pathway enrichment analysis for proteins with elevated Klac peptides highlighted in (**h**). (j) Schematic representation of HLA-B on the cell membrane, depicting its three extracellular domains (α1, α2 and α3) associated with β2-microglobulin to form the MHC class I complex, and highlighting the lactylated lysine at position 210 (K210) within the α3 domain. (k) PRM validation of Klac peptides gained through Oktoberfest rescoring from lactyl-NHS–labeled HeLa lysates (related to *Homo sapiens* in Fig. 2c). Left: Stacked bar chart showing gained peptides detected (red) or undetected (blue) by PRM assay using the ChemIntelligence Scope approach. Right: Ridge plot displaying the CV distribution of the detected Klac peptides, accompanied by a pie chart illustrating the proportion of peptides with CV < 30% versus ≥ 30%.

Using ChemIntelligence Scope, we monitored 22,388 Klac peptides guided by retention time and spectral data from the ChemIntelligence Library, and developed 21 multiplexed PRM assays via Skyline and PRM Conductor. In chemically lactylated HeLa lysates analyzed on Stellar MS, these assays detected 17,494 precursors (**Fig. 3c, Supplementary Table 4**) with exceptional reproducibility—over two experiments performed five months apart, 15,656 and 15,841 peptides were detected, of which 13,966 overlapped (**Fig. 3c**). Retention-time and intensity profiles remained highly consistent across replicates (**Fig. 3d**). Notably, each PRM injection required only 200 ng of protein input, in sharp contrast to the mg-scale quantities typically needed for conventional antibody-based enrichment methods.

To rapidly assess the feasibility of ChemIntelligence Scope for querying endogenous Klac profiles in biosamples, we consolidated singe PRM assay for POC analysis by applying the targeted assay to lactate-treated HeLa cell lysates. Candidate peptides included in the consolidated assay were prioritized by abundance while meeting PRM Conductor criteria (absolute area > 500, signal-to-background ratio (S/B) > 3, relative abundance > 0.05, time correlation > 0.8, and > 3 high-quality transitions) and balancing retention-time distribution^17,18^. This process selected 1,214 Klac peptides, of which 822 Klac peptides exhibited a coefficient of variation (CV) < 30% after triplicate ChemIntelligence Scope profiling (**Supplementary Fig. S9a**, **Table 5**). Remarkably, only 2.07% of these peptides were detectable by affinity enrichment, underscoring the orthogonal and expanded coverage achieved by ChemIntelligence Scope (**Fig. 3e**). Motif analysis revealed distinct sequence preferences: ChemIntelligence Scope favored acidic-adjacent lysines (e.g., glutamate), whereas antibody enrichment preferentially captured lysine clusters (**Fig. 3f**). GO-CC analysis further highlighted broader subcellular coverage—exosomal and cytoplasmic proteins alongside nuclear hits—providing access to Klac substrates missed by antibody workflows (**Fig. 3g, Supplementary Fig. S9b-d**).

Applied the consolidated PRM assay to lactate-treated versus control HeLa cells at as low as 200 ng protein inputs, ChemIntelligence Scope quantified 43 significantly altered Klac peptides (40 upregulated, 3 downregulated), whereas antibody enrichment detected only one **(Fig. 3h**, **Supplementary Table 5**). KEGG pathway analysis of the 40 upregulated peptide-associated proteins highlighted two notable surface targets: integrin αV (ITGAV) and human leukocyte antigen B (HLA-B). They are enriched in regulation of the actin cytoskeleton and antigen processing/presentation, respectively (**Fig. 3i**). ITGAV—implicated in tumor progression, angiogenesis and metastasis across multiple cancer types^33^—was identified here as lactylated for the first time, suggesting a potential role for Klac in integrin-mediated cancer signaling. ChemIntelligence Scope also revealed a previously unreported lactylation site at HLA-B K210, located within the α3 domain that directly contacts CD8 on cytotoxic T cells^34^, indicating a possible role in modulating T-cell activation and antitumor immunity (**Fig. 3j**).

Finally, we leveraged ChemIntelligence Scope to validate rescoring-gained peptides from our Prosit–Oktoberfest workflow. Five PRM injections targeting 4,156 rescoring-rescued Klac peptides detected 3,686 hits in lactyl-NHS–labeled HeLa samples, with 2,260 meeting CV < 30% criteria (**Fig. 3k**, **Supplementary Table 6**), confirming the authenticity of our deep learning-enhanced identifications and their readiness for targeted PRM assays. In summary, ChemIntelligence Scope enables scalable, reproducible, antibody-free mapping of the lactylproteome from low-input samples (200 ng per injection), expands target space beyond antibody-based enrichment methods, and connects discovery directly to quantitative validation—broadening both the technical reach and biological insight into Klac regulation.

### Ultra-low-input ChemIntelligence Scope achieves single-cell Klac detection

Having established that hundreds of Klac peptides can be detected from lactate-treated HeLa cells using 200 ng of protein input, we next evaluated ChemIntelligence Scope performance at ultra-low input, down to a single cell. To minimize losses and maximize signal, we employed a single-column LC configuration without a trap-elute setup, coupled with FAIMS to reduce chemical noise and matrix interference^35^. Using this optimized setup, we reconstructed the Klac ChemIntelligence Library comprising 18,012 Klac peptides (**Supplementary Table 7**).

Because single-cell experiments allow only one injection, we developed a consolidated PRM assay targeting 1,224 Klac peptides across 1,003 proteins within a 30-min gradient based on the reconstructed library for single-cell ChemIntelligence Scope analysis (**Fig. 4a, Supplementary Table 8 and Fig. S10a**). The selected Klac substrates span diverse pathways (**Supplementary Fig. S10b**). Klac profiling was then conducted across a wide input range—from bulk cell samples (10^3^ to 10^5^ cells) prepared by serial dilution to isolated cell samples (1, 5, 10, 20, 50 and 100 cells) obtained via cellenONE sorting. As expected, signal intensity decreased (**Fig. 4b-c**) and CV increased with declining input (**Fig. 4d-e, Supplementary Table 8**), with individual peptides showing consistent attenuation across all levels (**Fig. 4c**).

**Figure 4.**
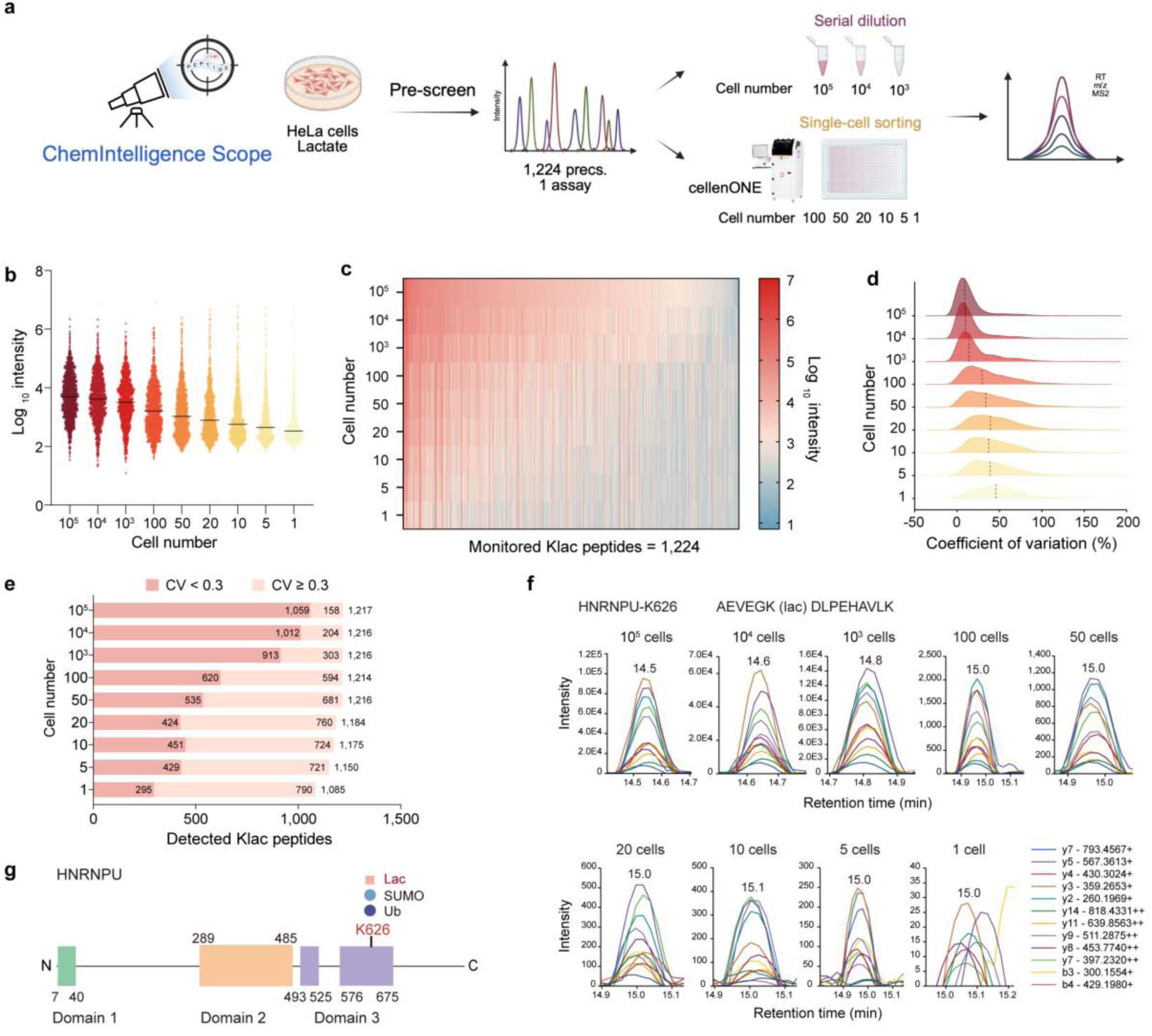
ChemIntelligence Scope enables enrichment-free quantification of the lactylproteome down to single-cell level. (a) Schematic workflow of high-throughput, high-sensitivity lactylproteome profiling from ultra-low-input samples using ChemIntelligence Scope. (b) Violin plots showing the distribution of detected Klac peptide intensities in samples ranging from single-cell to 10^5^-cell inputs. (c) Heatmap of monitored peptide intensities ranging from single-cell to 10^5^-cell samples. Peptides are ordered by their average intensity in the 10^5^-cell group. Undetected peptides are colored white in heatmap. (d) Ridge plots of CV distributions across input levels from single-cell to 10^5^-cell samples. Dashed lines indicate the median CV for each input level. (e) Stacked bar charts showing peptide CV distributions across input levels with CV < 30% in dark pink and CV ≥ 30% in light pink: numbers within bars indicate detected peptides. (f) XICs of targeted precursor-fragment ion transitions for the Klac peptides bearing HNRNPU-K626 across input levels, with annotated XICs shown on the right. This peptide was selected for being in the top 25% of CVs within the 100-cell group. (g) Domain organization of HNRNPU derived from AlphaFold predictions (AF-Q00839-F1-v4), with the K626 site highlighted. This residue has been previously reported to undergo SUMOylation and ubiquitination, and is identified here as lactylated.

Remarkably, at single-cell resolution 1,085 peptides were detected, including 295 with CV < 30% (**Fig. 4e**), demonstrating robust detection at the extreme sensitivity limit. As an example, AEVEGK(lac)DLPEHAVLK—bearing Klac at HNRNPU-K626 and falling at the 25th percentile of CV values for the 100-cell group—showed a clear input-dependent increase in extracted ion chromatograms (XICs) of its fragment ions (**Fig. 4f**), with signals remaining readily discernible even at the single-cell level (**Supplementary Fig. S10c**). Notably, K626 has previously been reported to undergo SUMOylation ^36^ and ubiquitination^37^, making it as a PTM hotspot and suggesting PTM crosstalk with significant functional implications (**Fig. 4g**). Taken together, these observations position ChemIntelligence Scope as a dedicated platform for Klac analysis, enabling quantification down to the single-cell level with sensitivity and resolution unattainable by conventional affinity-enrichment approaches.

### Decoding lactylation dynamics in scarce clinical specimens

After establishing that ChemIntelligence Scope sensitively profiles Klac in ultra-low-input cell line samples, we next asked whether this approach could meet the challenge of quantifying Klac peptides from scarce patient-derived tissues and primary cells—sample types central to addressing key biological questions but notoriously limited in material.

To address such analytical demands, we implemented a strategic two-step workflow: a pre-screening stage using sample-type-matched population cells to prioritize high-abundance Klac peptides, followed by targeted PRM analysis of the clinical specimens (**Fig. 5a**). For example, to enable Klac profiling in non-small-cell lung cancer (NSCLC) biopsies (**Supplementary Table 9**), we first screened lactylated H1299 cells—a representative NSCLC cell line—and refined the PRM assay to 1,628 targeted precursors using the same criteria as bulk cells. In this optimized set, 1,255 Klac peptides were reproducibly quantified (CV < 30%) across triplicates (**Fig. 5b**, **Supplementary Table 10**). Leveraging these H1299-prioritized PRM assays, ChemIntelligence Scope quantified 890 unique Klac peptides (1,115 sites across 593 proteins) from three paired NSCLC tumor and tumor-adjacent tissue sections (**Fig. 5c**, **Supplementary Fig. S11a**, **Tables 9** and **11**), consuming only ∼200 ng of digested protein per assay in three consecutive runs per sample. This capacity—to detect ∼1,000 Klac peptides from a single 10-μm-thick NSCLC section—stands in sharp contrast to prior antibody-based methods, which fall short in low-input contexts. Moreover, the platform can be readily extended to proteome-wide Klac mapping by pooling multiple sections with the ChemIntelligence Library–based PRM assays.

**Figure 5.**
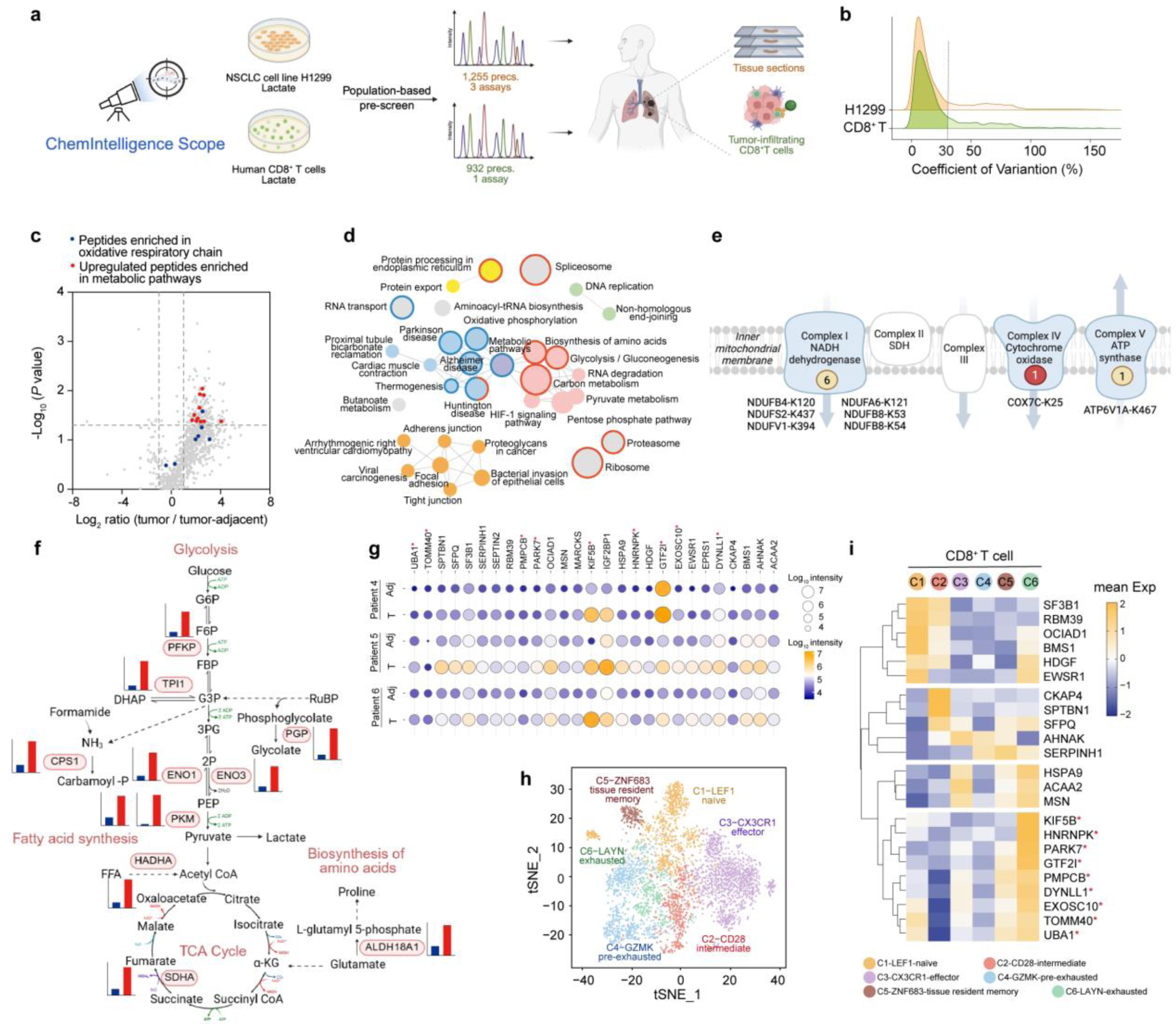
Quantitative mapping of dynamic lactylation changes in low-input clinical specimens using ChemIntelligence Scope. (a) Optimized ChemIntelligence Scope workflow with population-based pre-screening step to prioritize high-abundance targets for PRM analysis in low-input clinical samples. (b) Ridge plots showing CV distributions of Klac peptides from triplicate ChemIntelligence Scope PRM assays in H1299 cells and in *in vitro*-expanded CD8⁺ T cell lysates. Peptides with CV < 30% (dashed line) were deemed quantifiable. (c) Volcano plot of Klac peptide in tumor versus adjacent tissue. Significantly altered peptides were defined as tumor/tumor-adjacent ratio > 2 or < 0.5, *P* value < 0.05 by ratio paired *t*-test (n=3 biological replicates/group). Peptides from proteins enriched in the oxidative respiratory chain are highlighted in blue, and upregulated peptides from proteins enriched in metabolic pathways are highlighted in red. (d) KEGG pathways enriched for proteins with quantified Klac peptides (Fisher’s exact test, with Benjamini-Hochberg adjusted). Pathways uniquely detected by ChemIntelligence Scope are framed in blue and those enriched in tumors are framed in red. (e) Newly identified Klac sites (absent from **Supplementary Table 3**) mapped onto mitochondrial respiratory complexes, related to **Supplementary Fig. S11g**. Circles mark ChemIntelligence Scope-detected sites, colored by regulation status (red, upregulated; yellow, unchanged) and labeled with site positions. (f) Elevated lactylation on enzymes in glycolysis, TCA cycle, fatty acid, and amino acid biosynthesis, with accompanying bar plots showing intensities of the corresponding Klac peptides. (g) Bubble plot of significantly upregulated Klac peptides (tumor/tumor-adjacent Ratio > 2, *P* value < 0.05 by ratio paired *t*-test) in tumor-infiltrating CD8⁺ T cells. Peptides marked with red asterisks correspond to proteins whose encoding genes are highly expressed in cell cluster 6 (C6) in panel (**h**). (h) t-SNE projection of 3,335 single CD8⁺ T cells from 11 LUAD patients, revealing six major clusters defined by prominent marker genes: C1 (LEF1⁺, naive), C2 (CD28⁺, intermediate), C3 (CX3CR1⁺, effector), C4 (GZMK⁺, pre-exhausted), C5 (ZNF683⁺, tissue-resident memory), and C6 (LAYN⁺, exhausted). Points represent individual cells colored by cluster identity. (i) Heatmap of normalized expression of genes encoding proteins corresponding to upregulated Klac peptides summarized in (**g**) across the six CD8^+^ T cell clusters shown in (**h**). Red asterisks denote genes enriched in the exhausted cluster C6.

Functional annotation of the quantified peptides by GO and KEGG revealed enrichment in metabolic pathways (**Fig. 5d**, **Supplementary Fig. S11b–d**). Importantly, by targeting a pre-defined list rather than relying on relative affinity to Klac-specific antibodies, ChemIntelligence Scope captured Klac sites on low-abundance proteins that are often missed by enrichment-based approaches (**Supplementary Fig. S11e**), including 651 previously unreported sites (**Supplementary Fig. S11f**) on essential enzymes, such as subunits of mitochondrial complexes I, IV, and V (**Fig. 5e**, **Supplementary Fig. S11g–h**).

Of the quantified peptides, 163 were significantly upregulated in tumors (**Fig. 5c**), in line with the overall Klac increase observed by immunoblotting (**Supplementary Fig. S12a**). Notably, many of these Klac changes occurred independently of total protein abundance (**Supplementary Fig. S12b**), pointing to potential site-specific regulatory roles. Enrichment analysis linked altered Klac sites to key metabolic enzymes in carbon metabolism, glycolysis/gluconeogenesis, and amino acid biosynthesis (**Fig. 5d** and **5f, Supplementary Fig. S12c**). Structural mapping reinforced their functional associations, exemplified by K14 lactylation at the active site of triosephosphate isomerase 1 (TPI1) (**Supplementary Fig. S12d–e**).

We next explored Klac biology in patient tumor-infiltrating T cells, a compelling focus for tumor-microenvironment research given lactate’s reported role in immune regulation. Using the same pre-screening strategy, *in vitro*-expanded CD8⁺ T cells from healthy donors were profiled to develop a prioritized PRM assay comprising 932 validated Klac peptide precursors (CV < 30%) (**Fig. 5a–b, Supplementary Table 12**). Applying ChemIntelligence Scope to CD8⁺ T cells isolated from three paired lung adenocarcinoma (LUAD) tumor and adjacent tissue specimens yielded 484 Klac peptides across 433 proteins (**Supplementary Fig. S13a–b**, **Tables 9** and **13**). Functional enrichment analyses across biological process, molecular function, and cellular component categories highlighted diverse targets (**Supplementary Fig. S13c**). Consistent with the NSCLC findings, over 60% of Klac sites detected by ChemIntelligence Scope were missed by affinity-enrichment methods (**Supplementary Fig. S13d**). Among the ChemIntelligence Scope–detected peptides, 27 peptides carrying 30 Klac sites mapped to 27 proteins were significantly upregulated in tumor-infiltrating CD8⁺ T cells compared with CD8⁺ T cells isolated from adjacent tissues (**Fig. 5g**). Analysis of published LUAD single-cell RNA-seq data^38^ identified an exhausted CD8⁺ T cell cluster (C6) (**Fig. 5h**, **Supplementary Fig. S14**), within which nine genes corresponding to proteins with upregulated Klac levels in **Fig. 5g** were selectively enriched (**Fig. 5i**). This correlation supports a potential regulatory role of Klac in modulating proteins associated with T cell exhaustion.

### Extension of ChemIntelligence Scope to antibody-free mapping of nicotinylation

Having demonstrated the capability of ChemIntelligence Scope for antibody-independent profiling of Klac, we next examined whether such ChemIntelligence Library–based strategy could be generalized to broad lysine acylations analysis for low-input samples. Because the library is synthesized through similar NHS–ester chemistry targeting proteinaceous lysine, the ChemIntelligence Scope framework can be readily adapted to map other acyl-modifications.

To showcase this versatility, we extended ChemIntelligence Scope to Knic (**Fig. 6a**)—a nascent, nicotinate-induced PTM that links metabolic status to epigenetic and transcriptional regulation^28^. The lack of commercially available antibodies has so far hindered its proteomic characterization. Building on previous Klac-exemplified ChemIntelligence workflow, we synthesized nicotinyl-NHS (**Supplementary Fig. S15**) and optimized its chemical induction in HeLa lysates (**Supplementary Fig. S16**), followed by in-depth proteomic analysis using an Orbitrap Astral mass spectrometer (**Fig. 6a**). The CycIm and linear immonium (LinIm) ions characteristic of acyl modifications^25,30^ were readily observed for the chemically generated Knic peptides (**Fig. 6b**). Analogous to Klac, the Knic ChemIntelligence Library was completed within two days, comprising 26,605 Knic peptides corresponding to 3,273 proteins and 22,624 modified sites (**Fig. 6c**, **Supplementary Table 14**). Using this library, we established 24 PRM assays covering 16,759 Knic peptides (**Supplementary Table 15**).

**Figure 6.**
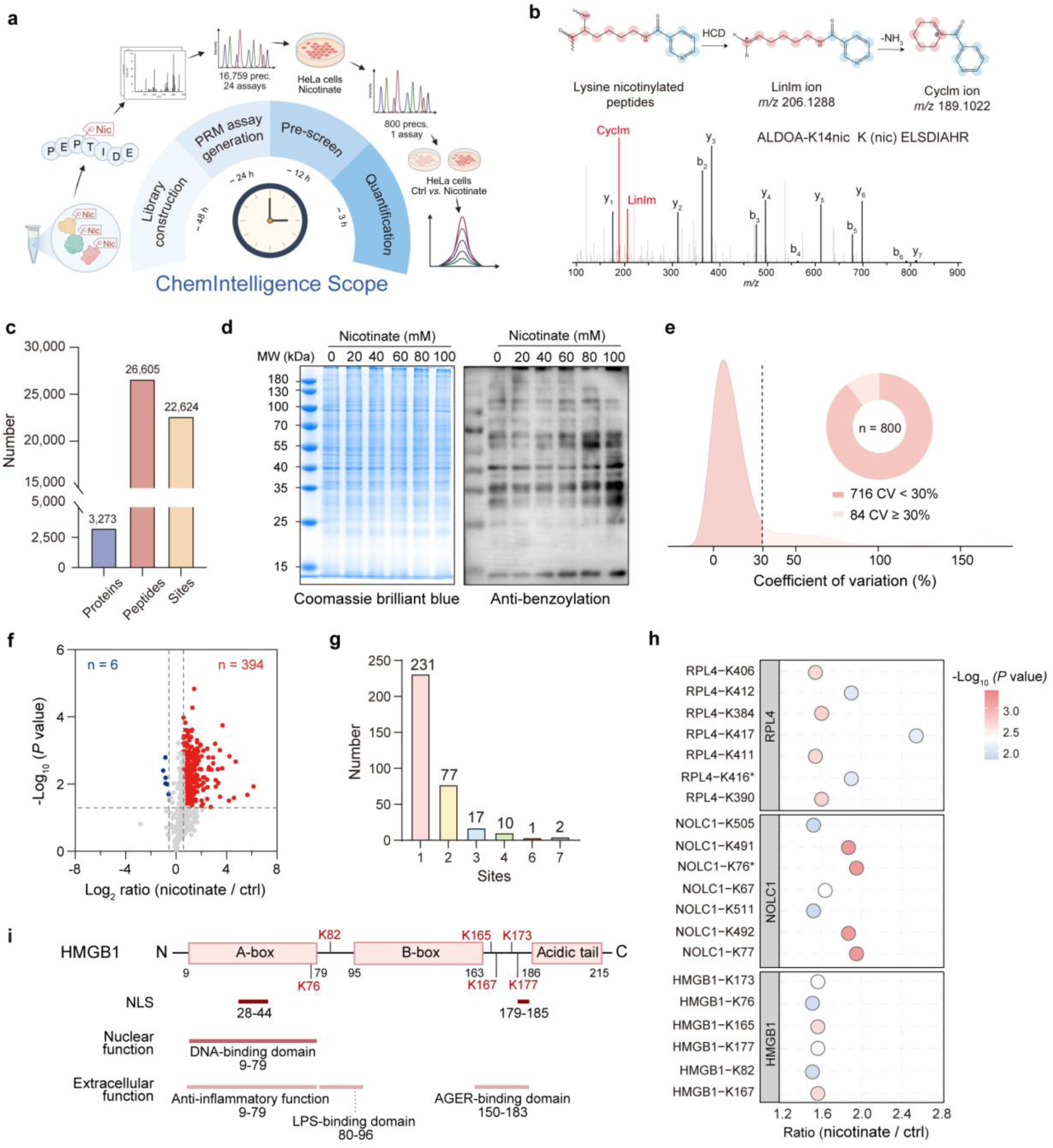
Expansion of ChemIntelligence Scope to Knic enables rapid, quantitative Knic profiling without the need to develop specific antibodies. (a) Schematic workflow of ChemIntelligence Scope for nicotinylation profiling, including library construction (∼48 h), PRM assay generation (∼24 h), pre-screening (∼12 h), and quantification (∼3 h). The entire confirmation and quantification process can be completed within 5 days. (b) Formation pathways of the characteristic CycIm and LinIm ions from Knic peptides. Pink circles represent the lysine-derived atoms and blue circles represent the nicotinic acid–derived atoms. The bottom panel shows a representative spectrum of a Knic peptide from the Knic ChemIntelligence Library with annotated CycIm and LinIm ions. (c) Bar plot showing the numbers of identified Knic proteins, peptides and sites in Knic ChemIntelligence Library. (d) Immunoblot validation of Knic upregulation in nicotinate–treated HeLa cells using an anti-benzoylation antibody previously reported to cross-react with Knic in the absence of a commercial antibody. (e) Ridge plot showing CV distribution of Knic peptides from triplicate ChemIntelligence Scope PRM analyses in nicotinate–treated HeLa cells. Peptides with CV < 30% (to the left of the dashed line) were considered quantifiable, and the accompanying pie chart shows the proportion of peptides with CV < 30% versus ≥ 30%. (f) Volcano plot of Knic peptides quantified by ChemIntelligence Scope, illustrating nicotinate treatment-associated changes in HeLa cells. Peptides meeting significance criteria (nicotinate/control Ratio > 1.5 or < 0.67, *P* value < 0.05 by ratio paired *t*-test, n=3 biological replicates/group) are highlighted in red (upregulated) and blue (downregulated), respectively. (g) Bar chart depicting the number of modified Knic sites per protein among the upregulated peptides identified in panel (**f**). (h) Bubble plot illustrating the upregulation trends of Knic peptides in panel (**f**), highlighting the three proteins with the greatest number of upregulated peptides. Asterisks indicate sites detected in multiple peptide precursors, with only the highest-intensity precursor shown. (i) Domain architecture of HMGB1, with lysines exhibiting upregulated Knic levels highlighted in red. Key functional regions, including the NLS and domains linked to nuclear or extracellular functions, are indicated.

We next set out to map the Knic landscape using a targeted assay. Knic levels were induced in HeLa cells by nicotinate treatment—a condition reported to elevate Knic—and confirmed by immunoblotting with a benzoylation-specific antibody known to cross-react with Knic^39^ (**Fig. 6d**). Following the same processing workflow as ChemIntelligence Scope for Klac, serial PRM assays were performed on nicotinate-treated lysates, and Knic peptides were prioritized into a consolidated PRM set comprising 800 peptides, 927 modified sites, and 548 proteins. Triplicate analyses via ChemIntelligence Scope reproducibly quantified the targeted peptides, with 716 showing CV < 30% (**Fig. 6e**, **Supplementary Table 16**). Consistent with immunoblotting, 394 peptides across 338 proteins were significantly increased after nicotinate treatment (**Fig. 6f**). Notably, each PRM run required only ∼30 min, enabling additional serial assays to further expand coverage of the largely unexplored Knic proteome.

Overall, the complete workflow—from a new acylation discovery to proteome-level quantification—was accomplished within 5 days, underscoring an efficient, entirely antibody-free paradigm for rapid acylation characterization. Frequency analysis revealed that roughly one-third of the proteins with upregulated Knic peptides contained multiple modification sites, indicating a widespread modification with potential substrate preference (**Fig. 6g**). For example, 60S ribosomal protein L4 (RPL4) and nucleolar and coiled-body phosphoprotein 1 (NOLC1) each carry seven sites, while high mobility group box 1 (HMGB1) harbors six (**Fig. 6h**); all three proteins have been previously linked to oxidative stress responses. HMGB1, a prototypical damage-associated molecular pattern (DAMP) protein, possesses multiple PTMs known to modulate inflammatory and immune signaling^40^, suggesting that nicotinate—beyond its role as an NAD⁺ precursor in energy metabolism and redox homeostasis^41^—may fine-tune oxidative stress-associated pathways via direct protein nicotinylation. The 6 upregulated Knic sites span DNA-binding (K76), lipopolysaccharide-binding (K82), and advanced glycosylation end product-specific receptor (AGER)-binding (K165–K177) regions (**Fig. 6i**), the latter critical for NF-κB activation^40,42^. The nicotinylated K177’s proximity to the nuclear localization signal further implies a role in regulating HMGB1 shuttling and stress-triggered release^40^, pointing to Knic as a modulator of downstream inflammatory signaling.

## DISCUSSION

In this study, we developed an acyl-NHS ester–based ChemIntelligence strategy to construct peptide reference library for two representative acyl-lysine PTMs, Klac and Knic. To our knowledge, this collection represents the largest LC–MS/MS spectral resource for lactylated (> 256,000 peptides) and nicotinylated (> 26,000 peptides) sequences, each annotated with iRT values and high-resolution MS and MS/MS spectra. Unlike conventional synthetic peptide libraries, our workflow circumvents labor-intensive peptide synthesis by exploiting rapid, low-cost (∼10 USD) chemical derivatization to generate extensive acylated peptide repertoires. For Klac, this approach yielded 22,388 confidently identified peptides from only three DDA runs.

The library’s authenticity and breadth were rigorously validated through diagnostic ion analysis, comparison with antibody-enriched datasets, and validation of representative sequences using synthetic standards. These evaluations demonstrated that the library accurately reflects endogenous lactyl- and nicotinyl-proteomes, offering large-scale, unbiased coverage across proteins with diverse functions and subcellular localizations. Because the same derivatization chemistry can be tailored to other acyl-lysine species, the ChemIntelligence workflow establishes a generalizable blueprint for building comprehensive acyl-proteome reference resources beyond Klac and Knic.

Then, we demonstrated broad utilities of ChemIntelligence Library in three application areas. First, in computational development, we trained two Prosit-Lac models to predict iRTs and fragment ion intensities, then integrated them into the Oktoberfest rescoring workflow—analogous to how deep learning approaches such as Prosit-Cit^43^ have advanced peptide identification. This integration markedly increased peptide identifications across diverse unenriched and affinity-enriched lactylproteome datasets from human and other species. Additional Klac peptides detected in HeLa lysates were validated by PRM and incorporated into the ChemIntelligence Scope–monitoring panel, with the framework demonstrating high efficiency, cross-species scalability, and robust performance in complex PTM contexts.

Second, to overcome the long-standing reliance on antibodies for acyl-proteome mapping, we developed ChemIntelligence Scope—a targeted serial-PRM platform powered by the ChemIntelligence Library. ChemIntelligence Scope reproducibly maps Klac peptides from as little as ∼200 ng of peptide input down to true single-cells, enabling antibody-free interrogation of low-input samples. Applied to frozen tumor sections and tumor-infiltrating CD8⁺ T cells from patient biopsies, ChemIntelligence Scope generated patient-specific Klac profiles and uncovered potential regulatory roles in T-cell exhaustion—insights previously inaccessible to conventional workflows. By deliberately targeting biologically relevant proteins via library-defined PRM assays, we anticipate ChemIntelligence Scope will become a powerful, scalable tool for routine acyl-proteome analysis in rare yet functionally important cell subsets across both clinical and fundamental biological studies.

We further extended ChemIntelligence Scope’s sensitivity to the frontier of single-cell proteomics, consistently detecting over 300 Klac peptides without isobaric-carrier multiplexing. This coverage far exceeds state-of-the-art low-input acetyl-proteomics approaches, such as Iseq-Kac^44^ (5 Kac peptides from 100 cells) and SCoPE^45^ (< 10 acetyl peptides from single cell). Together, these computational and experimental advances position ChemIntelligence Scope as a benchmark technology for global lactylproteome analysis, unlocking high-resolution Klac landscapes in rare and heterogeneous cell populations.

Third, we leveraged the library for rapid characterization of a novel PTM, Knic. Conventional discovery pipelines require time-intensive development of modification-specific antibodies—often 3∼4 months of work with high risk of failure, particularly for small, unstable, or immune-tolerant moieties. Even successful antibodies may exhibit cross-reactivity or sequence bias, missing many true sites. In contrast, the ChemIntelligence Library enables direct queries against modified peptides of interest, independent of antibody availability or binding preference. Leveraging the ChemIntelligence Scope workflow, we achieved Knic discovery and proteome-level mapping within just 5 days—a dramatic acceleration compared with antibody-based approaches—and demonstrated both the speed and flexibility of library-driven PTM exploration.

However, limitations of the current approach should be acknowledged. First, unlike fully synthetic libraries that provide explicit sequence identities, our method infers sequences via database searching and FDR control from unpurified Klac and Knic peptide mixtures, making accuracy dependent on search algorithms. Second, synthetic libraries often exploit multiple fragmentation methods (HCD, CID, ETD) and diverse instruments, whereas our study used only HCD acquisitions, with Prosit-Lac trained solely on Orbitrap and Astral datasets. Expanding to additional fragmentation modes, instrument platforms, and computational training sets will increase the generality of the framework.

In summary, the ChemIntelligence Library is a scalable, low-cost spectral resource for Klac and Knic, readily extensible to other acyl-lysine modifications such as crotonylation, succinylation, and butyrylation. Its utility spans Klac-specific Prosit-Lac model training to enhance peptide identifications across species, the creation of ChemIntelligence Scope for precise, antibody-free PRM profiling from immune cells, tumor biopsies, and true single-cell inputs, and the rapid characterization of Knic compared with the months required for antibody-based workflows. By uniting the breadth, efficiency of chemical-library generation with the precision of targeted PRM and the ability to interrogate low-input samples, this platform sets a benchmark for acyl-proteome analysis beyond Klac and Knic, accelerating mechanistic discovery and expanding acyl-proteome mapping from fundamental biology to translational and clinical research.

## METHODS

### Human participants and ethical approval

The protocol for this study was approved by the ethics committees of The First Affiliated Hospital with Nanjing Medical University (2022-SR-760). Whole blood for CD8^+^ T cell isolation was obtained from three healthy volunteers. Tumor tissues and tumor-adjacent lung tissues were obtained from six non-small-cell lung cancer (NSCLC) patients who underwent surgical resection (**Supplementary Table 9**). This study was conducted in accordance with the principles of Declaration of Helsinki.

### Cell lines and chemicals

HeLa, H1299, MC38, CHO and NRK-49F cells were purchased from American Type Culture Collection (ATCC) and cultured at 37 ℃ in a 5% CO_2_ atmosphere. HeLa, MC38, CHO and NRK-49F cells were cultured in Dulbecco’s Modified Eagle Medium (DMEM), and H1299 cells were cultured in RPMI-1640 medium. All culture media were purchased from Gibco (Thermo Fisher Scientific, USA) and supplemented with 10% fetal bovine serum (FBS, Excell, cat. no. FSP500), 100 U/mL penicillin and 1 μg/mL streptomycin (Thermo Fisher Scientific, cat. no. 15070-063). LC-MS grade water and acetonitrile (ACN) were obtained from Merck. All chemicals were purchased from Sigma-Aldrich unless otherwise specified. *Escherichia coli* and *Saccharomyces cerevisiae* were purchased from Sangon Biotech (Shanghai, China) and cultured in Luria–Bertani (LB) broth at 37 ℃ with shaking at 220 rpm for 16 h.

### Human CD8^+^ T cell isolation and culture

#### Peripheral CD8⁺ T Cells

Human peripheral blood was collected from healthy, non-fasted individuals into heparin-coated tubes (Konsfi, cat. no. 231112). Peripheral blood mononuclear cells (PBMCs) were isolated using Lymphoprep (STEMCELL Technologies, cat. no. 18061) through density gradient centrifugation at 800 g for 20 min at 25 ℃ with the brake off. Residual red blood cells were subsequently lysed with red blood cell lysis buffer (RBC lysis buffer, Beyotime, cat. no. C3702). CD8^+^ T cells were further separated using the EasySep Human CD8^+^ T cell Isolation Kit (STEMCELL Technologies, cat. no. 17953). Freshly purified CD8^+^ T cells were activated with 2.5 μg/mL anti-human CD3 and 2.5 μg/mL anti-human CD28 antibodies (BioLegend, cat. no. 317326 and 302934), cultured with ImmunoCult-XF T Cell Expansion Medium (STEMCELL Technologies, cat. no. 10981) supplemented with 100 U/mL penicillin, 1 μg/mL streptomycin and 10 ng/mL recombinant human interleukin-2 (IL-2; PeproTech, cat. no. 200-02).

#### Tumor-infiltrating CD8^+^ T cells

Fresh tumor and tumor-adjacent tissues from lung adenocarcinoma (LUAD) patients were washed with ice-cold phosphate-buffered saline (PBS), weighed, and mechanically dissociated into small pieces. The tissues were then digested with 4 mL RPMI-1640 medium supplemented with 2% FBS containing 1 mg/mL collagenase IV (Gibco, cat. no. 17104019) and 0.1 mg/mL Deoxyribonuclease I (Sigma-Aldrich, cat. no. DN25). Following a 30-min digestion at 37 ℃ on a shaking platform, tissue suspensions were sequentially filtered through a 70-μm cell strainer, centrifuged at 300 g for 10 min at 4 ℃, and lysed using RBC lysis buffer to lyse the residual red blood cells. The resulting cell pellets were used for CD8^+^ T cell separation using the REAlease CD8 (TIL) MicroBead Kit (Miltenyi Biotec, cat. no. 130-096-495) according to the manufacturer’s instructions.

### Synthesis and characterization of lactyl-NHS and nicotinyl-NHS

#### Synthesis of lactyl-NHS and nicotinyl-NHS

Lactic acid (180 mg, 1.0 equiv) or nicotinic acid (246.2 mg, 1.0 equiv) and *N*-hydroxysuccinimide (278.2 mg, 1.2 equiv) were dissolved in anhydrous acetonitrile and stirred at room temperature for 15 min. *N*, *N*’-Dicyclohexylcarbodiimide (494.4 mg, 1.2 equiv) was suspended in ACN, then added dropwise to the reaction mix. The solution was stirred at room temperature for 6 h. After filtration, the filtrate was concentrated under reduced pressure, then resuspended in dry hexane and stirred at room temperature for 12 h to induce precipitation. The precipitate was collected and concentrated to yield (2,5-dioxopyrrolidin-1-yl) (2*S*)-2-hydroxypropanoate (lactyl-NHS) or 2,5-dioxopyrrolidin-1-yl nicotinate (nicotinyl-NHS), respectively.

#### Characterization of lactyl-NHS and nicotinyl-NHS

Lactyl-HNS was measured using the 1290 Infinity II UHPLC system (Agilent Technologies, Palo Alto, CA, USA) coupled with the 6546 quadrupole time-of-flight (Q-TOF) mass spectrometer (Agilent). The chromatographic separation was performed on an XBridge Amide column (4.6 × 100 mm, 3.5 μm, Waters). The mobile phase consisted of solvent A (10 mM ammonium acetate solution, pH 9.0) and solvent B (ACN). The gradient was set as follows: 0-1.0 min, 85% B; 1.0-12.0 min, 85%-30% B; 12.0-14.0 min, 30% B; 14.0-14.5 min, 30%-85% B; 14.5-26.0 min, 85% B. The flow rate was set at 0.4 mL/min and the column temperature was set as 40 ℃. Following separation, the Agilent 6546 Q-TOF was operated in the negative mode (ESI-) for lactyl-NHS detection. The following parameters were used: liquid nebulizer, 45 psi; nitrogen drying gas, 8 L/min; drying gas temperature, 320; ESI capillary voltage, 3500 V; MS scan, *m/z* 50-1100; MS/MS scan, *m/z* 25-850; fragmentor voltage, 80 V.

For structural confirmation of lactyl-HNS and nicotinyl-NHS, ^1^H NMR and ^13^C NMR spectra were acquired (500 MHz for ^1^H and 125 MHz for ^13^C) on Avance III HD 500 MHz NMR spectrometer (Bruker BioSpin GMBH, Ettlingen, Germany). Samples were dissolved in deuterated dimethyl sulfoxide (DMSO-*d*_6_) with a concentration of approximately 20 mg/mL. Chemical shifts were referenced to the residual solvent signal. For ^1^H NMR of lactyl-NHS, the following resonances were observed: δ 6.05 (d, *J* = 5.7 Hz, 1H), 4.57 (p, *J* = 6.7 Hz, 1H), 2.84 (s, 4H), and 1.43 (d, *J* = 6.9 Hz, 3H). For ^13^C NMR of lactyl-NHS, the following signals were observed: δ 171.21, 170.64, 64.93, 25.98, and 21.10. For ^1^H NMR of nicotinyl-NHS, the following resonances were observed: δ 9.25 (s, 1H), 9.00 (d, J = 4.8 Hz, 1H), 8.49 (d, J = 8.0 Hz, 1H), 7.75 – 7.70 (m, 1H), 2.94 (s, 4H). For ^13^C NMR of nicotinyl-NHS, the following signals were observed: δ 170.63, 161.49, 156.19, 150.93, 138.28, 125.07, 121.59, 40.40, 26.08.

### Expression and purification of recombinant proteins

The his-tagged plasmid pET22b-ALDOA, pET22b-ENO1 and pET22b-CBR1 were transfected into BL21(DE3) cells by heat shock. Single colonies were selected and then grown at 37 ℃ with shaking at 220 rpm in LB broth until the OD_600_ reached 0.8. Isopropyl-*β*-D-thiogalactopyranoside (IPTG) was then added to a final concentration of 0.2% to induce protein expression. The resulting mixtures were centrifuged, and the precipitates were resuspended in lysis buffer containing 5 mM imidazole, 150 mM NaCl, 50 mM tris(hydroxymethyl)aminomethane (Tris), 1 mg/mL lysozyme and 1% protease and phosphatase inhibitor cocktails (ApexBio, cat. no. K1007 and K1013). The suspensions were then incubated on ice for 10 min and subsequently sonicated. The lysates were centrifuged, and the supernatants were collected and loaded onto the Ni-NTA prepacked chromatographic columns (Sangon Biotech, cat. no. C600792-0105) for enrichment of his-tagged proteins. The concentrated ALDOA, ENO1 and CBR1 were eluted with imidazole buffer followed by filtration through the 3 kDa ultrafiltration column (Merck, cat. no. UFC9003) to obtain the purified recombinant proteins.

### Chemical labeling with lactyl-NHS or nicotinyl-NHS and proteomic sample preparation

#### Peptide

The reaction mixture (50 μL) comprised 200 mM HEPES (pH 8.0) in 50% ACN, 5 mM lactyl-NHS dissolved in ACN and 1 μM synthetic peptide DGADFAKWR. The mixture was kept on a shaker at 25 ℃ for 1.5 h and quenched with 4 μL 5% hydroxylamine at room temperature for 15 min, acidified to pH 2-3 with trifluoroacetic acid (TFA) and evaporated to dryness. Samples were desalted with C_18_ ZipTips (Millipore), evaporated to dryness and stored at -80 ℃ prior to analysis.

#### Recombinant proteins

Recombinant proteins were diluted in a buffer containing 10 M urea (Sigma-Aldrich, cat. no. U5378) and 200 mM HEPES (pH 8.0, Beyotime, cat. no. ST092) to a final volume of 30 µL, resulting in a working solution of 0.67 µg/µL protein with a final urea concentration greater than 6 M. The diluted proteins were reduced with 10 mM tris (2-carboxyethyl) phosphine (TCEP, Sigma-Aldrich, cat. no. 75259) at 56 ℃ for 30 min, followed by alkylation with 20 mM Iodoacetamide (IAM, Sigma-Aldrich, cat. no. I1149) at 25 ℃ for 20 min in the dark. Lactyl-NHS was then added to reach a final concentration of 1 mM after optimization. The mixture was kept on a shaker at 25 ℃ for 1.5 h and quenched by 2.5 μL 5% hydroxylamine at room temperature for 15 min. Subsequently, methanol, chloroform and water were added to the lysate at a ratio of 4:1:3:1 by volume to precipitate proteins. Precipitated proteins were washed twice with cold methanol, re-solubilized in 8 M urea containing 200 mM HEPES, and digested with LysC (1:100, *w/w*, Promega, cat. no. VA1170) at 25 ℃ for 4 h. The solution was diluted to achieve a final urea concentration less than 1 M, followed by trypsin digestion (1:100, *w/w*, Promega, cat. no. V5111) at 37 ℃ overnight. Peptides were acidified with 0.1% formic acid (FA, Thermo Fisher Scientific, cat. no. 28905), desalted using C_18_ ZipTips, evaporated to dryness and stored at -80 ℃.

#### Cell lysate

Cells were harvested, washed three times with ice-cold PBS and lysed in 8 M urea containing 200 mM HEPES (pH 8.0) with protease and phosphatase inhibitor cocktails. Lysates were sonicated and centrifuged at 16,000 g for 10 min at 4 ℃, and the supernatants were collected. Protein concentrations were determined by bicinchoninic acid (BCA) assay. Lysates were then reduced with 10 mM TCEP at 56 ℃ for 30 min and alkylated by 20 mM IAM at 25 ℃ for 20 min in dark, labeled with 10 mM lactyl-NHS (after optimization) or 15 mM nicotinyl-NHS at 25 ℃ for 1.5 h on a shaker. Subsequently, cell lysates were processed for precipitation, re-solubilization, digestion and desalting as described in the section of recombinant protein level.

#### Tissue lysate

Longissimus dorsi muscles of *Sus scrofa*, *Bos taurus* and *Ovis aries* were dissected, weighed, and finely minced. The tissue pieces were washed five times with PBS until no visible coloration remained. Samples were then lysed in 1% (w/v) sodium dodecyl sulfate (SDS) solution and homogenized at 4 ℃ for 10 min at 60 Hz. After centrifugation, the insoluble material was discarded, and the supernatant was collected. Proteins were precipitated, resuspended in 200 mM HEPES (pH 8.0) containing 1% SDS, and diluted to about 5 µg/µL. Subsequent labeling and downstream processing were performed following the same protocol as for cell line samples.

### High-pH reversed-phase HPLC fractionation

Fractionation was used to construct the expanded Klac ChemIntelligence Library. Briefly, digested lactyl-NHS–labeled cell lysates were dissolved in HPLC phase A buffer (5% ACN containing 10 mM ammonium formate) and fractionated on a Prominence LC-20AD system (Shimadzu Corporation, Kyoto, Japan) using a ZORBAX 300Extend-C_18_ (4.6 mm × 250 mm, 5 μm, Agilent) at 35 ℃. Phase B consisted of 20% ACN in 10 mM ammonium formate buffer. The gradient was set as follows: 0-5 min, 5% B; 5-79 min, 5-50% B; 79-81 min, 50–100% B; 81-98 min, 100% B; 98-120 min, 100-5% B. The flow rate was set as 0.6 mL/min. The effluent was collected every 1.5 min. For 10-fraction samples, every 10 fractions were pooled; for 24-fraction samples, fractions from each 24-fraction cycle were combined across cycles; 48-fraction samples were analyzed without pooling. The lyophilized fractions were dissolved in 60 µL of 0.1% FA aqueous buffer, desalted with C_18_ ZipTips, evaporated to dryness and stored at -80 ℃ before analysis.

### Proteomic sample preparation of standard-input HeLa cells

To induce lactylation for pre-screening experiments, HeLa cells were treated with 100 mM sodium lactate, H1299 cells with 80 mM sodium lactate, and human CD8⁺ T cells with 40 mM sodium lactate, all for 48 h. For pre-screening of nicotinylation, HeLa cells were treated with 100 mM sodium nicotinate. In parallel, for quantitative analyses in HeLa cells, both sodium lactate–treated and sodium nicotinate–treated groups were prepared together with their corresponding untreated controls cultured under identical conditions. After treatment, cells were washed three times with ice-cold PBS, harvested, and lysed in 8 M urea containing 50 mM ammonium bicarbonate (Sigma-Aldrich, cat. no. 09830) solution with protease and phosphatase inhibitor cocktails. Lysates were sonicated and centrifuged to collect supernatants. Protein concentrations were determined by BCA assay. Lysates were then reduced with 10 mM dithiothreitol (DTT, Sigma-Aldrich, cat. no. 43815) at 56 ℃ for 30 min and subsequently alkylated with 20 mM IAM at 25 ℃ for 20 min in the dark. Subsequently, cell lysates were processed for digestion and desalting as described above.

### Proteomic sample preparation with serial dilution

HeLa cells treated with 100 mM sodium lactate for 48 h were washed twice with PBS and digested with 0.25% trypsin-EDTA buffer (Gibco, cat. no. C25200-072) at 37 ℃ for 4 min to obtain a single-cell suspension. Cells were counted using Automated Cell Counter C100 (RWD Life Science Co, Shenzhen, China) and diluted accordingly. Suspensions containing 10^3^, 10^4^ and 10^5^ cells were aliquoted and centrifuged at 5,000 rpm for 5 min, and the supernatant was carefully removed. The cells were resuspended in 100 μL lysis buffer (pH 8.5) containing 1% (*w/v*) sodium deoxycholate (SDC, Aladdin, cat. no. S104198), 10 mM TCEP, 40 mM 2-chloroacetamide (CAA, Sigma-Aldrich, cat. no. 22790) and 100 mM Tris-HCl. The lysates were then boiled at 95 ℃ for 5 min^46^, homogenized, and centrifuged at 14,000 g for 5 min. Subsequently, the supernatant was collected and digested overnight at 37 ℃. Proteolysis was quenched by adding FA to a final concentration of 0.5%, followed by centrifugation at 16,000 g to remove SDC. The supernatant was then desalted using C_18_ ZipTips, evaporated to dryness, and stored at -80 ℃ until proteomic analysis.

### Proteomic sample preparation with cellenONE

HeLa cells treated with 100 mM sodium lactate for 48 h were washed twice with PBS and digested with 0.25% trypsin-EDTA at 37 ℃ for 4 min to obtain a single-cell suspension. Cells were diluted with PBS to 100–200 cells per µL in a 200-µL tube and dispensed by the cellenONE (Cellenion SASU, Berlin, Germany) into an Eppendorf 384-well twin.tec PCR LoBind plate (Eppendorf SE, Hamburg, Germany) preloaded with 1 µl of master mix buffer (consisting of 0.2% DDM (D4641-500MG, Sigma-Aldrich), 100 mM TEAB, 20 ng/µl trypsin) per well. Cell morphology, density, and diameter were assessed by the instrument, and only cells within a 15–35 µm size range and with an elongation factor ≤ 1.8 were selected for sorting. Depending on experimental design, wells were loaded with either single cell or defined groups of 5, 10, 20, 50, or 100 cells, with each cell co-deposited with 400 pL of PBS. Following sorting, the plate underwent controlled incubation at 37 ℃ with 70% relative humidity for 2 h. After the reaction, the temperature was reduced to 20 ℃ to stabilize the samples. Subsequently, 3.5 µL of 0.1% TFA was added to each well. The plate was sealed and transferred to the Vanquish Neo UHPLC system, enabling direct injection of 4.5 µL per sample for downstream MS analysis.

### Affinity enrichment for lactylated peptides

Endogenous cell samples were generated by treating cells with sodium lactate for 48 h (HeLa, 100 mM; H1299, 80 mM; expanded human CD8⁺ T cells, MC38, CHO, and NRK-49F, 40 mM). Basal (unstimulated) counterparts were processed in parallel. Cell pellets were lysed and digested following the standard-input cell protocol described above. For tissue samples, longissimus dorsi muscle from pig, cow, and sheep—processed with or without lactyl-NHS labeling—was lysed and digested as described above. Protein A/G agarose beads (Santa Cruz Biotechnology, cat. no. sc-2003) were washed with NETN buffer (0.5% NP40, 1 mM EDTA, 100 mM NaCl, 50 mM Tris-HCl, pH 8.0) and blocked with 5% bovine serum albumin (BSA). The beads were then conjugated with the anti-lactylation antibody (Micron Bio, cat. no. WM101) via overnight incubation at 4 ℃. Trypsin-digested peptides were incubated with the antibody-conjugated beads for 5 h at 4 ℃ with rotary shaking. Beads were washed three times with NETN buffer, and lactylated peptides were eluted with 0.1% TFA. Eluted peptides were evaporated, desalted, and stored at -80 ℃ before proteomic analysis.

### Proteomic sample preparation of clinical samples

For tissue sections, paired tumor and tumor-adjacent tissues from three NSCLC patients were embedded in optimal cutting temperature (OCT) compound (Dakewe, cat. no. 4081511). Serial sections were prepared using a Cryostat (Dakewe, cat. no. CT520): 50-μm-thick for immunoblotting and 10-μm-thick for proteomics. The 10-μm-thick sections were washed with PBS and resuspended in 100 μL lysis buffer (pH 8.5) containing 1% (*w/v*) SDC, 10 mM TCEP, 40 mM CAA and 100 mM Tris-HCl. For tumor-infiltrating CD8^+^ T cells, purified CD8^+^ T cells were similarly processed. The lysates were then digested and desalted as described for serially diluted samples.

### Proteomic analysis by LC-MS/MS

#### Orbitrap Eclipse Tribrid analysis

The 10-fraction and 24-fraction digested lactyl-NHS–labeled cell lysates were analyzed by an Orbitrap Eclipse Tribrid mass spectrometer equipped with an EASY-nano LC 1200 LC system (Thermo Fisher Scientific, MA, USA). The 10-fraction digested lactyl-NHS–labeled cell lysates were analyzed in duplicate. Chromatographic separation of digested lactyl-NHS–labeled cell lysates was performed on an Acclaim PepMap RSLC column (75 μm × 250 mm, Thermo Fisher Scientific) at 50 ℃ using a flow rate of 300 nL/min with mobile phase A (0.1% formic acid in water) and mobile phase B (ACN/H_2_O, 8:2, *v/v*), under a 90-min gradient. The digested recombinant proteins were separated using the same column condition with a 60-min gradient. The mass spectrometer was operated in a data-dependent acquisition (DDA) mode. MS1 spectra were acquired in the *m/z* range of 350-1,800 at 120,000 resolution on the Orbitrap with a maximum automated gain control (AGC) of 4e^5^. Isolation window for precursors was set as 1.6 Th. For MS2 acquisition, fragmentation was conducted by higher-energy collisional dissociation (HCD) with the Normalized Collision Energy (NCE) set at 32%. MS2 spectra were obtained with the first mass set at *m/z* 100 at 30,000 resolution, and a maximum AGC of 5e^4^.

#### Orbitrap Astral analysis

The digested lactyl-NHS–labeled lysates (non-fractionated and 24 fractions) and the affinity-enriched lactylated peptides from HeLa cells were analyzed using DDA mode on an Orbitrap Astral mass spectrometer (Thermo Fisher Scientific, USA) coupled with a Vanquish Neo UHPLC system (Thermo Fisher Scientific, USA) using “Trap elute” mode. The digested non-fractionated lactyl-NHS–labeled lysates were analyzed in triplicate using DDA mode to construct the proof-of-concept (POC) Klac ChemIntelligence Library. Mobile phase A consisted of 0.1% FA in water, and phase B of ACN-H_2_O (8:2, *v/v*) with 0.1% FA. Peptides was first loaded on a PepMap Neo trap cartridge (300 μm × 5 mm, 5 μm, Thermo Fisher Scientific) and then separated on an Aurora Elite column (75 μm × 150 mm, 1.7 μm, IonOpticks) using a 30-min gradient at 50 ℃ and a flow rate of 300 nL/min. MS1 spectra were acquired in the *m/z* range of 300-1,600 at 180,000 resolution on the Orbitrap with a 300% normalized AGC target. Isolation window for precursors was set as 1.6 Th. MS2 spectra were obtained by HCD fragmentation using 30% NCE with a scan range of *m/z* 100-2,000 and a 200% normalized AGC target.

Label-free quantification (LFQ) of proteins in paired tumor and tumor-adjacent tissue sections from three NSCLC patients was conducted in data-independent acquisition (DIA) mode. MS1 spectra were acquired in the *m/z* range of 400-1,000 at 240,000 resolution with a 500% normalized AGC target and 5e^6^ absolute AGC target, using a 5-ms maximum injection time. Data were acquired using 300 consecutive precursor isolation windows of 2 Th width, spanning an *m/z* range of 400-1,000. Isolated ions were fragmented by HCD with 30% NCE. MS2 spectra were acquired in the *m/z* range of 100-2,000, with a 500% normalized AGC target, 5e^4^ absolute AGC target, and a maximum injection time of 3 ms.

#### Orbitrap Astral Zoom analysis

The cross-species affinity-enriched lactylproteome data, the digested non-fractionated nicotinyl-NHS–labeled lysates were analyzed using DDA mode on an Orbitrap Astral Zoom mass spectrometer (Thermo Fisher Scientific, USA) coupled with a Vanquish Neo UHPLC system (Thermo Fisher Scientific, USA) using “Trap-elute” mode. All settings were the same as on Orbitrap Astral with the addition of “Pre-accumulation” on.

The digested non-fractionated lactyl-NHS–labeled lysates from HeLa cells were analyzed using DDA mode on an Orbitrap Astral Zoom mass spectrometer coupled with a Vanquish Neo UHPLC system using “Direct injection” mode to create the ultra-low-input Klac ChemIntelligence Library. Orbitrap Astral Zoom mass spectrometer was connected to the FAIMS pro interface at compensation voltage −48 V. The other settings were the same as those used in “Trap-elute” mode.

#### timsTOF HT analysis

The 48-fraction lactyl-NHS–labeled lysate digests were analyzed on a timsTOF HT mass spectrometer (Bruker Daltonics, Bremen, Germany) coupled with a NanoElute2 LC system (Bruker Daltonics, Bremen, Germany). Mobile phases A (0.1% FA in water) and B (0.1% FA in ACN) were delivered at 300 nL/min. Peptides were separated using a 60-min gradient on Aurora Ultimate C_18_ columns (75 µm × 150 mm and 250 mm, 1.7 μm, IonOpticks) at 50 ℃. The mass spectrometer operated in DIA parallel accumulation-serial fragmentation (PASEF) mode (10 scans/cycle). The trapped ion mobility spectrometry (TIMS) system was configured with a 166-ms accumulation/ramp. Scan ranges spanned *m*/*z* 100-1,700 for the mass and 1/*K*_0_ 0.75-1.50 V*s/cm^2^ for ion mobility. Suitable precursor ions for PASEF-MS/MS were selected in real time from the TIMS-MS survey scans based on *m*/*z*-ion mobility coordinates. Dynamic exclusion parameters: intensity threshold = 500, target intensity = 5,000, exclusion duration = 0.4 min. Quadrupole isolation widths were set to 2 Th for precursors with *m*/*z* < 700, ramped linearly from 2 Th to 3 Th for *m/z* between 700 and 800, and fixed at 3 Th for *m/z* > 800. CE was linearly ramped with ion mobility from 90 eV at 1.5 V*s/cm^2^ to 26 eV at 0.75 V*s/cm^2^.

#### Stellar analysis

All 200-ng loading amount PRM samples were analyzed on a Stellar mass spectrometer (MS) (Thermo Fisher Scientific, USA) coupled with a Vanquish Neo UHPLC system using “Trap-elute” mode. All liquid phase conditions were identical to those employed on the Orbitrap Astral instrument. Ultra-low-input samples were analyzed in “Direct injection” mode without a trap column and with the Orbitrap Astral mass spectrometer connected to the FAIMS Pro interface at a compensation voltage of −48 V. For calibrating retention time (RT), adaptive RT was activated. For each distinct sample condition—including changes in matrix, treatment, or formulation, an rtbin file was first generated before acquirement, and then applied to subsequent relative data acquirement. MS1 spectra were acquired in the *m/z* range of 300-1,600 at a rate of 33 kDa/s, with a cycle time of 1.14s. Maximum injection time mode was set to custom, which is equivalent to 10 ms. AGC target was set to standard with the absolute AGC target of 3e^4^. MS2 spectra were obtained by HCD fragmentation (NCE 30%) with a scan range of *m/z* 200-1,500 at a rate of 125 kDa/s. AGC target was set to standard with the absolute AGC target of 1e^4^, and the maximum injection time was set to dynamic. The loop control was set to “All”.

### Proteomic data analysis

#### Database searching for Klac ChemIntelligence Library

Proteomic data for Klac ChemIntelligence Library were searched against the UniProt human proteome (UniProt_Human_reviewed_15-05-2023.fasta) using SEQUEST in Proteome Discoverer 3.1 (Thermo Fisher Scientific) for POC and ultra-low-input Klac ChemIntelligence Library, and PEAKS Studio XPro (Bioinformatics Solutions) for the remaining data in the expanded Klac ChemIntelligence Library. The enzyme was set to trypsin, allowing a maximum of three missed cleavages and semi-specific digestion. Carbamidomethylation of cysteine was set as a fixed modification, while lactylation of lysine and oxidation of methionine were set as variable modifications, with a maximum of three variable modifications. A mass tolerance of 10 ppm was allowed for precursor ions and 0.02 Da for fragment ions for the data collected on the Orbitrap Astral, Orbitrap Astral Zoom and the Orbitrap Eclipse Tribrid. For the data acquired on the timsTOF HT, mass tolerances were set to 20 ppm for precursor ions and 0.04 Da for fragment ions. Peptide-spectrum matches (PSMs) were filtered to 1% false discovery rate (FDR) employing a target-decoy database search approach.

#### Database searching for Knic ChemIntelligence Library

Proteomic data for Knic ChemIntelligence Library were searched against the UniProt human proteome (UniProt_Human_reviewed_15-05-2023.fasta) using SEQUEST in Proteome Discoverer 3.1 (Thermo Fisher Scientific). The enzyme was set to trypsin, allowing a maximum of three missed cleavages and semi-specific digestion. Carbamidomethylation of cysteine was set as a fixed modification, while nicotinylation of lysine and oxidation of methionine were set as variable modifications, with a maximum of three variable modifications. A mass tolerance of 10 ppm was allowed for precursor ions and 0.02 Da for fragment ions. Peptide-spectrum matches (PSMs) were filtered to 1% false discovery rate (FDR) employing a target-decoy database search approach.

#### Database searching for DIA data

All DIA data were analyzed with Spectronaut 19 (Biognosys) using the library-free approach (directDIA+) and the same human reference database employed for DDA searches. The settings for digestion and modification were identical to those used in PEAKS Studio. Precursor filtering was set as *Q* value, and “Cross-run normalization” was enabled.

#### Database Searching for Prosit-Lac model building and Oktoberfest-rescoring

Raw files were searched using Fragpipe (v 22.0) with MSFragger (v4.1), IonQuant (v1.10.27), and Philosopher (v5.1.1). For Prosit-Lac model building, the UniProt *Homo Sapiens* reference proteome (reviewed, downloaded on 17.11.2023) was used. For Oktoberfest rescoring, species-specific UniProt reference proteomes were used, including: *Mus musculus* (reviewed, downloaded on 17.06.2025), *Sus scrofa* (unreviewed, downloaded on 22.07.2025), *Bos taurus* (unreviewed, downloaded on 22.07.2025), *Ovis aries* (unreviewed, downloaded on 07.07.2025), *Rattus norvegicus* (unreviewed, downloaded on 22.07.2025), *Cricetulus griseus* (unreviewed, downloaded on 22.07.2025), *Saccharomyces cerevisiae* (reviewed, downloaded on 20.12.2024), and *Escherichia coli* (reviewed, downloaded on 22.06.2023). Cysteine carbamidomethylation was specified as a fixed modification. Variable modifications included methionine oxidation and lactylation of lysine. Enzyme specificity was set to strict trypsin, allowing up to three missed cleavages. The precursor mass tolerance was set to 20 ppm, and MSFragger mass calibration was enabled. MSBooster rescoring was disabled. PSMs were filtered with Philosopher to a 1% FDR at the PSM level for standard searches.

#### Fragment ion intensity training data preparation and model training

To integrate the Klac ChemIntelligence Library with the PROSPECT dataset, we first performed a collision energy (CE) calibration using the Oktoberfest tool in combination with the Prosit 2020 HCD intensity models^26,47^ **(Supplementary Fig. S7a)**. For Orbitrap Eclipse Tribrid raw files, a single optimal CE value of 33 was identified, whereas Orbitrap Astral data showed four optimal CE values (28, 31, 32, and 33). Following calibration, raw files from the expanded Klac ChemIntelligence Library acquired on both the Orbitrap Tribrid Eclipse and Orbitrap Astral instruments were converted to the mzML format using the ThermoRawFileParser (https://github.com/compomics/ThermoRawFileParser). Each confidently identified spectrum by MSFragger was annotated with b- and y-ion coverage for fragment charges between 1 and 3. A mass tolerance of 20 ppm was applied to all annotations. Peptides shorter than 7 or longer than 30 amino acids, as well as spectra with precursor charges above 6, were excluded.

The intensity model was implemented in Python 3.10.0 with Keras 2.15.0 and TensorFlow 2.15.0. Its architecture, inspired by the Prosit 2020 TMT model^48^, includes sequence embedding layers, recurrent neural networks, and attention mechanisms designed to predict peptide fragmentation patterns. The sequence embedding was extended to include lactylation as an additional modification within the amino acid alphabet. Inputs to the model included the peptide sequence, normalized CE, precursor charge, and fragmentation method. To help the model account for differences between instruments, an extra input indicating the instrument type (Orbitrap Astral, Orbitrap Eclipse Tribrid, or Orbitrap Fusion Lumos Tribrid) was added **(Supplementary Fig. S7b)**. This instrument label was stored as an integer and converted into a 2-dimensional learned embedding, enabling the model to learn instrument-specific characteristics.

Annotated spectra from the Klac ChemIntelligence Library (∼2.3 million PSMs) were combined with the PROSPECT dataset^31^, resulting in ∼19.3 million PSMs. The combined dataset was split by unmodified sequences into 70% training, 20% validation, and 10% test sets to ensure no sequence overlap. The model was trained using the Adam optimizer with an initial learning rate of 0.0001, reduced by a factor of 0.8 if the validation loss did not improve over 4 epochs. Training was limited to 200 epochs but terminated after 182 epochs due to early stopping, which occurred when the validation loss failed to improve by at least 0.0001 for 8 consecutive epochs. The loss function was a trimmed version of the normalized spectral contrast loss^27^, excluding the worst-performing 1% of predictions per batch. Training was conducted using the DLOmix framework (https://github.com/wilhelm-lab/dlomix).

#### Retention time (iRT) training data preparation and model training

To calibrate iRT values across datasets, first, the iRTs of all unmodified peptides in the Klac ChemIntelligence Library were predicted using the Prosit 2019 iRT model^26^ and a representative RT for peptides in the Klac ChemIntelligence Library was selected using the highest-scoring spectrum for reference. Next, we fitted a linear regression between measured RTs and predicted iRTs of the unmodified peptides for each raw file separately. Last, these raw file-specific regression models were then used to transform the RTs of lactylated peptides into iRTs.

The iRT model was implemented with the same Python, Keras, and TensorFlow versions as the intensity model. Its architecture, inspired by the Prosit 2019 iRT model^26^, takes peptide sequences as input and predicts a single representative iRT value per peptide. The sequence embedding layer mirrors that of the intensity model, employing the same extended amino acid alphabet.

The dataset included ∼300,000 unmodified and lactylated peptides, which were split based on the unmodified sequences into 60% training, 20% validation, and 20% test sets, ensuring that no sequence was shared between the three sets. The model was trained using the Adam optimizer with an initial learning rate of 0.0001 for a maximum of 300 epochs, but training stopped early after 221 epochs due to early stopping, based on the same criteria as the intensity model. The loss function was a trimmed mean squared error (MSE), excluding the worst-performing 1% of predictions per batch. Training was also performed using the DLOmix framework.

#### Rescoring pipeline

Rescoring was performed as described in Oktoberfest^27^. Briefly, raw MS files and search engine results are processed by Oktoberfest to perform spectrum annotation, normalized collision energy (NCE) calibration, and indexed retention time (iRT) alignment **(Supplementary Fig. S7a)**. Peptide fragmentation intensity and iRT predictions from the Prosit-Lac models were retrieved via the Koina prediction service^32^ and used to compute an extensive set of features for false discovery rate (FDR) estimation using Percolator^49^.

#### Generation of target lists for PRM assay

The SEQUEST search results for data in the POC Klac ChemIntelligence Library were exported as pepXML files, and only peptides with a Percolator *q*-value < 0.01 were imported into Skyline (v24.1.1.398) to create a BiblioSpec library (Blib) as a reference for integrating PRM assays. Transition settings were adjusted, with peptide settings set to Trypsin [KR/P], maximum of 3 missed cleavages, and a 2-min retention time window (measured retention times used when available). Peptides of 6-45 amino acids were selected, with “Auto-select all matching peptides” enabled. Carbamidomethylation of cysteine was set as a fixed modification, and lactylation of lysine and oxidation of methionine as variable modifications (max 3 variable modifications and max 3 losses). Transition settings considered as +2 and +3 for precursor charges and +1 and +2 for ion charges, with an ion match tolerance of 20 ppm. DDA parameters were applied for full-scan settings. After peak picking from the raw data using the above parameters, any precursors lacking identified peaks were excluded. Subsequently, the PRM Conductor (v1.0.4344) in Skyline was configured as follows. For target refinement, “Keep All Precs.” option was selected. Method parameters were set with a minimum dwell time of 10 ms, an LC peak width of 10 s, and at least 7 points per peak, with a 1.4-min acquisition window. For each peptide, only a single charge state was retained. Finally, for the POC Klac ChemIntelligence Library, the PRM Conductor distributed 17,544 precursors across 18 assays. For ultra-low-input Klac ChemIntelligence Library, 18,157 precursors across 15 assays were distributed. For Knic ChemIntelligence Library, 16,811 precursors across 20 assays were distributed.

#### Construction of PRM assays

For all 3 ChemIntelligence Libraries, the target peptide lists were validated on a Stellar MS with a single injection per assay. The raw data were imported into Skyline for precursor filtration. Transition settings included peptides with precursor charges of +2 and +3, and fragment ions (b/y) with charges of +1 and +2. Ion match tolerance for the library was set to 0.5 Da, and 15 product ions were selected from the filtered ions. Instrument parameters were set with a 0.5 Da tolerance, and PRM was used for full-scan settings. After peak picking from the raw data using the above parameters, any precursors lacking identified peaks were excluded. Subsequently, the PRM Conductor was used to distribute the precursors across assays. The PRM Conductor was set with a minimum dwell time of 10 ms, an LC peak width of 9 s, and at least 8 points per peak, with a 1.4-min acquisition window. Finally, for the POC Klac ChemIntelligence Library, the PRM Conductor distributed 17,494 precursors across 21 assays. For ultra-low-input Klac ChemIntelligence Library, 18,012 precursors across 19 assays were validated. For Knic ChemIntelligence Library, 16,759 precursors across 24 assays were validated.

#### Pre-screening

For lactylation quantification, standard-input sodium lactate–treated HeLa cells were validated on 17,494 precursors across 21 single-injection assays on the Stellar MS. Using the same assays, sodium lactate–treated H1299 cells and CD8⁺ T cells were analyzed to support downstream Klac mapping. For ultra-low-input applications, standard-input sodium lactate–treated HeLa cells were assessed on the ultra-low-input Klac ChemIntelligence Library, covering 18,012 precursors across 19 assays. For nicotinylation quantification, sodium nicotinate–treated HeLa cells were validated on 16,759 precursors across 24 assays. All raw data were imported into Skyline. After peak picking using the same parameters as described in the construction of PRM assays section, any precursors lacking identified peaks were excluded. Subsequently, the PRM Conductor was used to filter high-quality precursors.

For precursor selection, standard-input and ultra-low-input lactylation samples as well as nicotinylation samples were filtered based on an absolute area > 500, a signal-to-background ratio (S/B) > 3, relative abundance > 0.05, time correlation > 0.8, and at least 3 good transitions. Method parameters in PRM Conductor were set with a minimum dwell time of 10 ms, an LC peak width of 9 s, and at least 8 points per peak, with a 1.4-min acquisition window. With “Balance Load” option activated, a single final assay was obtained for each sample type, containing 1,214 precursors for standard-input lactylation samples, 1,224 precursors for ultra-low-input lactylation samples, and 894 precursors for nicotinylation samples.

For clinical samples, two rounds of screening were carried out. In the first-round screening, precursors were retained if they met the following criteria: absolute area > 500, S/B > 3, relative abundance > 0.05, time correlation > 0.8, and at least 3 good transitions were retained. Method parameters in PRM Conductor were set with a minimum dwell time of 10 ms, an LC peak width of 9 s, and at least 8 points per peak, with a 1.4-min acquisition window. As a result, 1,628 lactylated precursors from H1299 cells were distributed across three PRM assays, and 2,050 lactylated precursors from CD8^+^ T cells were distributed across three PRM assays.

In the second-round screening for clinical samples, the three PRM assays were run in triplicates, and raw data were imported into Skyline using the same settings as those applied in the first-round screening. Any precursors lacking identifiable peaks were excluded, and those with a coefficient of variation (CV) greater than 30% were removed based on both Skyline analysis and manual inspection. Then, PRM Conductor was used to filter high-quality precursors. Precursors meeting an absolute area > 500, S/B > 3, relative abundance > 0.05, time correlation > 0.8 and at least 3 good transitions were retained. For H1299 cells, method parameters in PRM Conductor were set with a minimum dwell time of 15 ms, an LC peak width of 8 s, and at least 7 points per peak, with a 1.4-min acquisition window. Finally, 1,255 precursors without truncated peaks were retained, yielding three PRM assays for mapping the lactylproteome in tumor and tumor-adjacent tissue sections. For CD8^+^ T cells, method parameters in PRM Conductor were set with a minimum dwell time of 12 ms, an LC peak width of 8 s, and at least 7 points per peak, with a 1.4-min acquisition window. Under these settings, 1,710 precursors across three PRM assays were reproducibly quantifiable—defined as having no truncated peaks and CV < 30% across triplicate runs—for mapping the lactylproteome in tumor and adjacent-tumor infiltrating CD8^+^ T cells. In cases where infiltrating CD8⁺ T cells were present in trace amounts, the same method parameters were applied, with a 10 ms dwell time and “Balance Load” activated, yielding 932 precursors without truncated peaks in a single PRM assay. While the full three-assay design was used for samples with sufficient input, the compressed single-assay format ensured coverage in low-yield conditions.

### Quantification of lactylproteome and proteome from biological samples

#### Standard-input HeLa samples

Peptide concentrations were measured using a NanoDrop One spectrophotometer (Thermo Fisher Scientific, Wilmington, DE, USA) to ensure uniform sample loading of 200 ng prior to proteomic analysis. Raw mass spectrometry data were imported into Skyline using the same instrument parameters described above, with an ion match tolerance of 0.5 Da. Only peptides that were consistently detected in all paired control and sodium lactate–treated samples, showed no truncated peaks, and had a CV < 30% in the sodium lactate–treated group were considered for quantification. Lactylated peptides exhibiting a sodium lactate-to-control intensity fold change greater than 2 or less than 0.5 and a *P* value < 0.05 (two-sided ratio *t*-test) were considered significantly altered and retained for downstream analysis.

#### Ultra-low-input HeLa samples

For 1, 5, 10, 20, 50, 100 and 10^3^ cells, all samples were loading onto column for analysis. For 10^4^ and 10^5^ cells, peptide concentrations were measured to ensure uniform sample loading of 200 ng prior to proteomic analysis. Raw mass spectrometry data were imported into Skyline using the same instrument parameters described above, with an ion match tolerance of 0.5 Da. For quantification, TIC normalization was applied to all samples.

#### Clinical samples

Peptide concentrations were measured to ensure uniform sample loading of 200 ng prior to proteomic analysis. For tissue sections, one replicate was analyzed per assay across three independent PRM experiments. For infiltrating CD8⁺ T cell samples, patient 4 provided sufficient material for quantification across three assays, whereas patients 5 and 6 were analyzed with only a single PRM assay each, using “Balance Load” optimization.

Raw mass spectrometry data were imported into Skyline using the same instrument parameters described above, with an ion match tolerance of 0.4 Da. Only gene-specific, unique peptides detected consistently across all paired tumor and tumor-adjacent samples, without truncated peaks, were considered for quantification. Lactylated peptides exhibiting a tumor-to-adjacent-tissue intensity fold change greater than 2 or less than 0.5 and a *P* value < 0.05 (two-sided ratio *t*-test) were considered significantly altered and retained for downstream analysis.

For proteome-wide quantification in NSCLC tissue sections, statistical significance was determined by a two-sided ratio *t*-test with thresholds of fold change > 2 and *P* value < 0.05. Protein quantification for tumor-infiltrating CD8⁺ T cells was excluded due to insufficient sample quantity.

### Quantification of nicotinylproteome and proteome

Peptide concentrations were measured to ensure uniform sample loading of 200 ng prior to proteomic analysis. Raw mass spectrometry data were imported into Skyline using the same instrument parameters described above, with an ion match tolerance of 0.5 Da. Only peptides that were consistently detected in all paired control and sodium nicotinate–treated samples, showed no truncated peaks, and had a CV < 30% in the sodium nicotinate–treated group were considered for quantification. Nicotinylated peptides exhibiting a sodium nicotinate-to-control intensity fold change greater than 1.5 or less than 0.67 and a *P* value < 0.05 (two-sided ratio *t*-test) were considered significantly altered and retained for downstream analysis.

### Secondary structure analysis

Secondary structure prediction for lactylated sites in the expanded Klac ChemIntelligence Library and every lysine residue in lysine-containing proteins of the human proteome was performed using the AlphaFold 3.0 database. A custom Python script was used to extract the secondary structure for each lysine residue. The secondary structures were classified into five categories: α-helix, β-sheet, β-turn, loop, and unstructured.

### Protein class analysis

Protein class analysis was conducted as previously described^50^. Briefly, human proteins were systematically categorized into distinct functional classes using a hierarchical annotation framework. Custom python scripts were written to extract and process data from the Gene Ontology (GO) database. A library was constructed to place each GO term into a category (Transporter, channel and receptors; Enzymes; Gene expression and nucleic acid binding; Scaffolding, modulators and adaptors). Categories were assigned based on keyword matches (for example, “transporter” / “channel” for the first category, and “enzyme” / “kinase” for the second). Proteins with multiple qualifying GO terms were classified according to the above-mentioned order of categories. Those lacking relevant GO annotations were designated as “Unclassified”.

### Cosine similarity analysis

To estimate the spectral similarity between shared lactylated peptides identified by affinity-enrichment and those in the POC Klac ChemIntelligence Library, cosine similarity^51^ of the spectra derived from the same precursors was calculated using an in-house Python script. For each pair of mass spectra, fragment peaks were matched within a 0.01 Da tolerance window, followed by cosine similarity calculation of the matched peak intensity vectors:

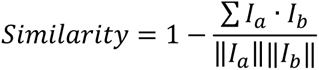

In this calculation, I_a_ and I_b_ represent the intensity vectors of matched peaks from the two spectra, and ||·|| denotes the L2 norm of the vectors. The calculation was implemented using the PyOpenMS toolkit, enabling efficient processing of large-scale mass spectrometry data.

### Immunoblotting

Briefly, 50-μm-thick tissue sections or nicotinate–treated cells were washed three times with ice-cold PBS and lysed in 60 μL RIPA buffer (New Cell & Molecular Biotech, cat. no. WB3100) supplemented with protease and phosphatase inhibitor cocktails. The lysates were homogenized and centrifuged at 14,000 g for 5 min at 4 ℃. Protein concentrations were determined by BCA assay. The lysates were diluted with 5× loading buffer (New Cell & Molecular Biotech, cat. no. WB2001), heated at 95 ℃ for 10 min, cooled. Proteins were separated by 10% sodium dodecyl sulfate-polyacrylamide gel electrophoresis (SDS-PAGE). For each sample, two identical gels were prepared in parallel; one gel was used for transfer onto polyvinylidene difluoride (PVDF) membranes for immunoblotting, and the other gel was subjected to Coomassie Brilliant Blue staining (Beyotime, cat. no. P0017) to assess protein loading. Proteins were then transferred onto polyvinylidene difluoride (PVDF) membranes, blocked with 5% nonfat dry milk in tris-buffered saline with 0.1% Tween 20 detergent (TBST) and incubated with primary antibodies at 4 ℃ overnight. The membranes were subsequently washed five times with TBST and incubated with horseradish peroxidase (HRP)-conjugated secondary antibody for 1 h at 37 ℃. The immunoblotted bands were captured on a ChemiDoc XRS+ system and analyzed by ImageLab software (Bio-Rad Laboratories, Hercules, CA, USA). The primary antibodies employed in this study include the antibodies against L-Lactylation Lysine (1:1000, Micron Bio, cat. no. WM-101), Benzoyllysine (1:1000, PTM Bio, cat. no. PTM-762) and β-actin (1:1000, Cell Signaling Technology, cat. no. 4967S).

### GO and KEGG pathway analysis

Lactylated proteins from the expanded Klac ChemIntelligence Library and the affinity-enriched lactylproteome were annotated with GO cellular component (CC) terms using the DAVID bioinformatics resource (https://david.ncifcrf.gov/). GO enrichment analysis of low-input sample data was performed using Metascape (http://metascape.org/), while KEGG pathway enrichment was carried out with KOBAS (http://bioinfo.org/kobas). Statistical significance for both GO and KEGG analysis was assessed using a one-sided Fisher’s exact test, with Benjamini-Hochberg-adjusted *P* < 0.05 as the threshold. Only gene sets containing at least 3 genes were included in the analysis.

### Single-cell RNA sequencing data processing

Single-cell RNA sequencing (scRNA-seq) data from 11 LUAD patients^38^ were reanalyzed using the corresponding gene expression matrices. Unsupervised clustering was performed using t-distributed stochastic neighbor embedding (t-SNE) and density peak clustering. The 2,000 genes with the highest standard deviation were selected, and principal component analysis (PCA) was applied to capture the major sources of biological variation. The top 30 principal components were subsequently used for t-SNE dimensionality reduction via the Seurat R package.

Based on characteristic gene expression signatures, CD8⁺ T cells were classified into six functional subtypes: C1-LEF1, C2-CD28, C3-CX3CR1, C4-GZMK, C5-ZNF683, and C6-LAYN. These subtypes exhibited distinct transcriptional profiles, reflecting heterogeneous functional states and differentiation trajectories. Subtype annotation was performed through hierarchical clustering and differential gene expression analysis across clusters. Marker genes were identified using the Wilcoxon rank-sum test, with an adjusted *P* value < 0.05 and an average Log₂ fold change > 0.25. Cell clusters were visualized using the DimPlot function in Seurat.

### Data availability

The Oktoberfest-rescored lactylproteome datasets were obtained from two public repositories. The label-free dataset OMIX006151 was obtained from OMIX (https://ngdc.cncb.ac.cn/omix/). In addition, three datasets were retrieved from the ProteomeXchange Consortium (http://proteomecentral.proteomexchange.org/), including PXD033991 (label-free), PXD051415 (SILAC), and PXD014870 (SILAC). **Supplementary Table 3** compiles all affinity-enriched lactylproteome data of *Homo sapiens* available from published literature as of May 2025. Experimental data used in this study and Oktoberfest-rescored data using Percolator have been deposited in the ProteomeXchange Consortium and can be accessed through the Consortium via the iProX partner repository^52^ with the dataset identifier PXD064554 and PXD063905. Source data are provided with this paper. The data used for training and testing both Prosit-Lac models are available on Hugging Face at https://huggingface.co/datasets/Wilhelmlab/Prosit-2025-lac-ms2 and https://huggingface.co/datasets/Wilhelmlab/Prosit-2025-lac-irt. Prosit-Lac models have been deposited on Zenodo (https://zenodo.org/records/17522962, https://zenodo.org/records/17523023).

### Code availability

Prosit-Lac models are available on Koina^32^ (https://koina.wilhelmlab.org/, https://github.com/wilhelm-lab/koina). Source code and scripts for model training are available on GitHub at https://github.com/wilhelm-lab/dlomix. Source code for rescoring using Oktoberfest are available on GitHub at https://github.com/wilhelm-lab/oktoberfest.

## ACKNOWLEDGEMENTS

This research was supported by the National Natural Science Foundation of China (grant 82322068 and 82173783 to H.Y., grant 82404703 to C.S.), the Natural Science Foundation of Jiangsu Province (BK20241590 to C.S.), the National Key Research and Development Program of China (2021YFA1301300 to H.H.), the Fundamental Research Funds for the Central Universities (2632022YC03 to H.Y.), the Overseas Expertise Introduction Project for Discipline Innovation (G20582017001 to H.H.), the Project of State Key Laboratory of Natural Medicines, China Pharmaceutical University (SKLNMZZ202402 to H.H.). This work was also supported by the Innovative Health Initiative Joint Undertaking (IHI JU) under grant agreement No 101165643. The JU receives support from the European Union’s Horizon Europe research and innovation programme and COCIR, EFPIA, EuropaBio, MedTech Europe and Vaccines Europe, EuropaBio and MedTech Europe and Breakthrough T1D and VeriSIM Life. Funded by the European Union, the private members, and those contributing partners of the IHI JU. Views and opinions expressed are however those of the author(s) only and do not necessarily reflect those of the aforementioned parties. Neither of the aforementioned parties can be held responsible for them. We thank Thermo Fisher Scientific China for providing the Orbitrap Astral and Stellar instruments, and Bruker China for the timsTOF HT instrument.

## AUTHOR CONTRIBUTION

H.Y., M.W., H.H. (Haiping Hao) and Z.G. conceived the project. C.S., Z.H., H.Y. and H.H. (Haiping Hao) designed the experiments. C.S., Q.Y., Z.H. and X.H. performed proteomic sample preparation and lactylation data mining. V.G. and M.W. trained the Prosit-Lac model and integrated it into Oktoberfest workflow. Z.H. and H.H. (Haoran Huang) conducted proteomic sample loading on the Orbitrap Astral and Stellar instruments. C.S., Q.Y. and Z.G. performed sample extraction from clinical tissues. Q.Y. and X.H. performed immunoblotting. C.S., Z.H., Q.Y., V.G., X.H., X.C. and Y.Z. performed data analysis. D.W. and Z.H. contributed to lactyl-NHS synthesis. Q.J. contributed to nicotinyl-NHS synthesis. C.S., Z.H. and Y.Z. conducted single-cell sorting. H.Y., C.S., Z.H. and V.G. wrote the manuscript.

