## Supplementary Information for "ChemIntelligence Enables Antibody-Free, Ultra-Low-Input Profiling of Lysine Lactylation and Diverse Acyl-Proteomes"

### Supplementary Figures

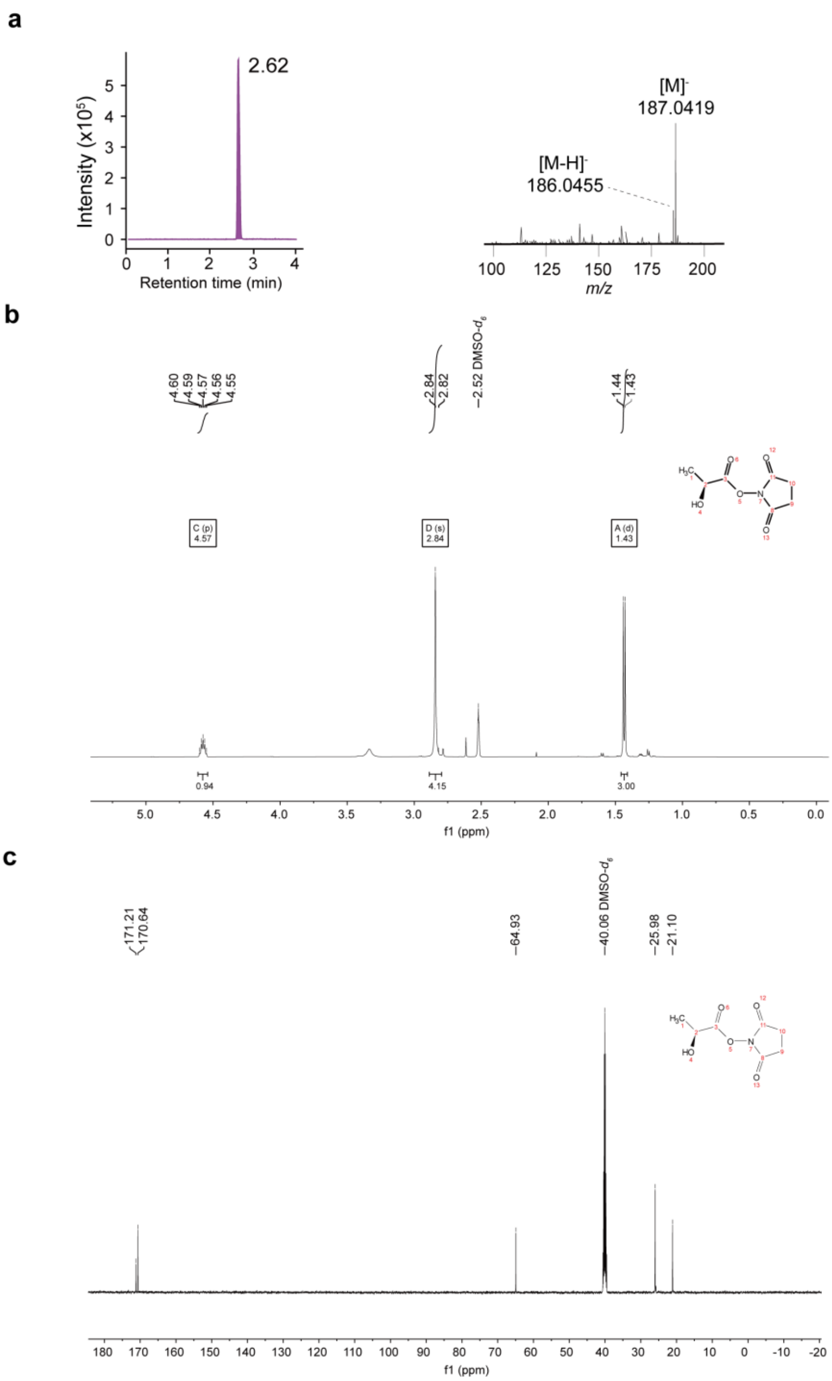


**Supplementary Figure 1. Synthesis and characterization of lactyl-NHS.**

(a) Extracted ion chromatogram (XIC, left) and MS spectrum (right) of lactyl-NHS.

(b) $^{1}$H NMR spectrum of lactyl-NHS in DMSO-$d_6$.

(c) $^{13}$C NMR spectrum of lactyl-NHS in DMSO-$d_6$.

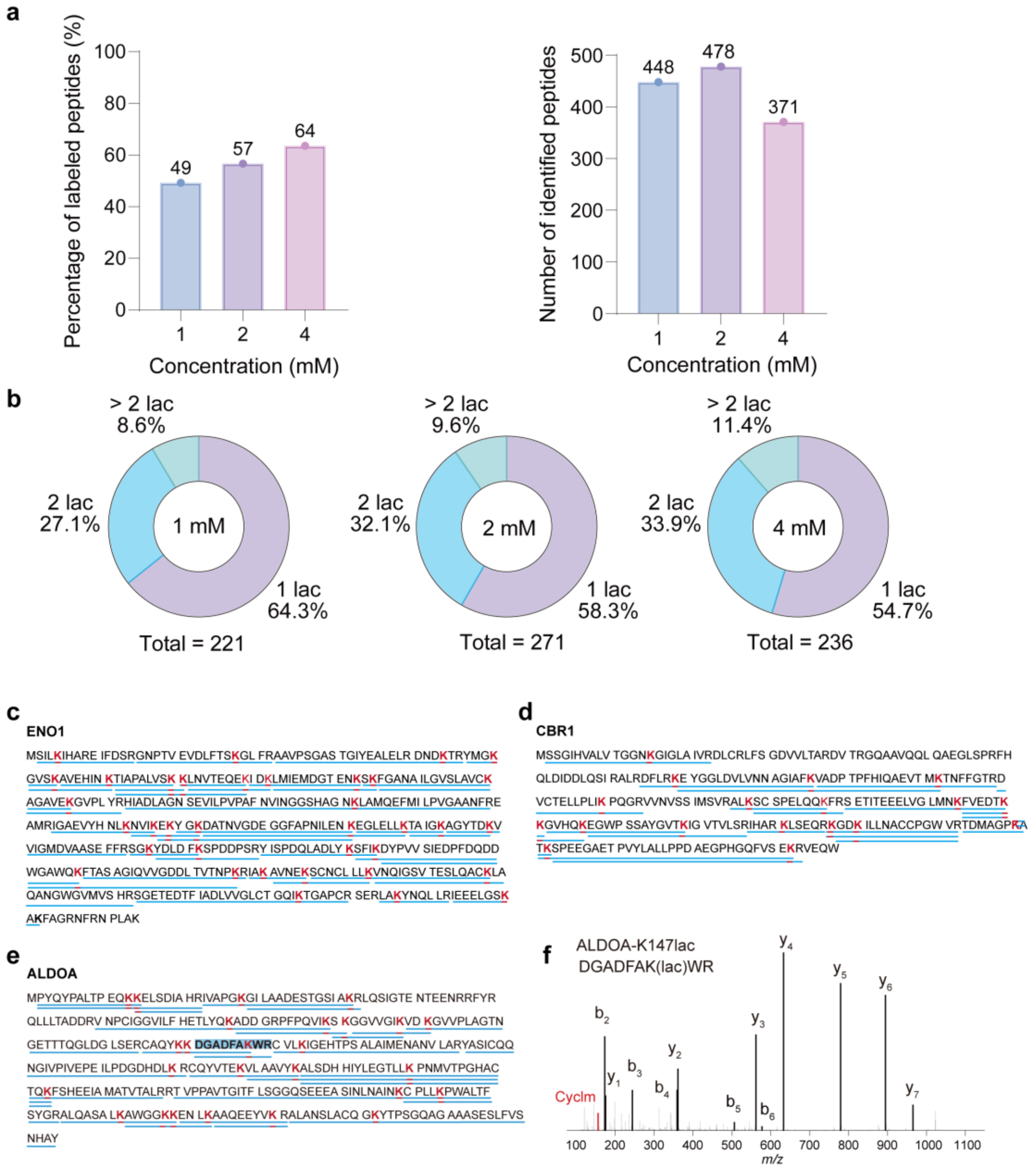


**Supplementary Figure 2. Reactivity of lactyl-NHS on recombinant proteins.**

(a) Percentage of lactyl-NHS–labeled peptides relative to the total identified peptides (left), and total peptide counts (right) in recombinant ALDOA at varying lactyl-NHS concentrations.

(b) Distributions of lactyl-NHS–labeled peptides with single (1 lac), double (2 lac), or multiple (> 2 lac) Klac sites in ALDOA at varying lactyl-NHS concentrations, showing reduced proportion of single-site labeling at higher concentrations.

(c–e) Lactylated lysines (red) in Klac peptides (blue) identified in lactyl-NHS–labeled ENO1 (c), CBR1 (d) and ALDOA (e), determined by Orbitrap Eclipse Tribrid.

(f) Representative MS/MS spectrum of a tryptic Klac peptide from lactylated ALDOA.

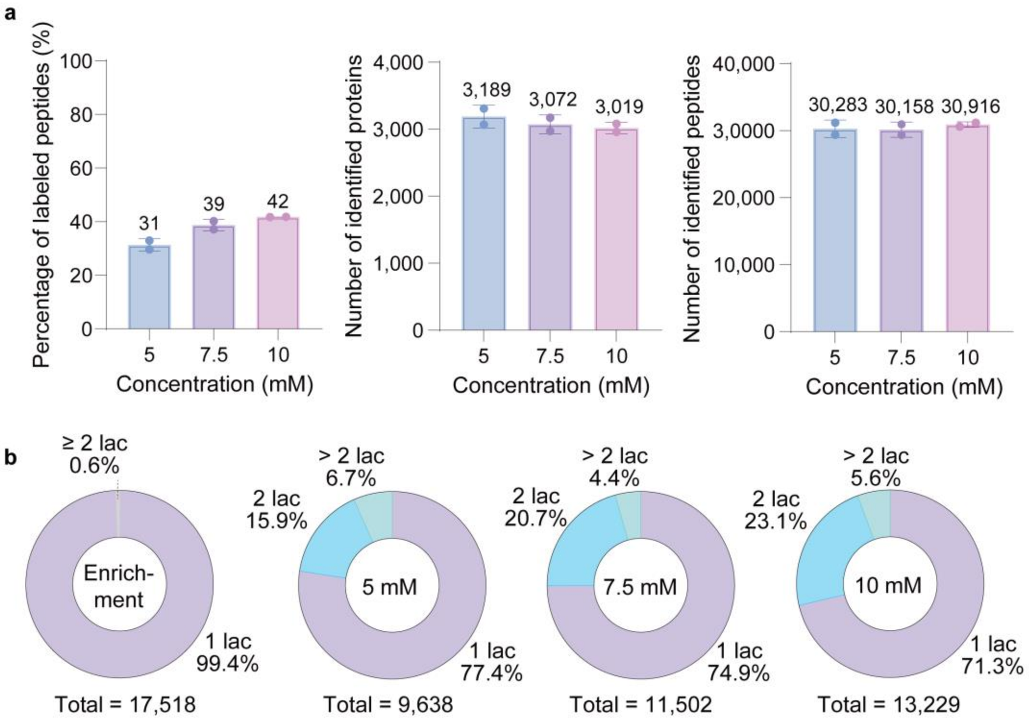


**Supplementary Figure 3. Reactivity of lactyl-NHS on cell lysates.**

(a) Percentage of lactyl-NHS–labeled peptides relative to the total identified peptides (left), total identified proteins (middle) and total peptides (right) in CD8$^+$ T cell lysates at the indicated lactyl-NHS concentrations.

(b) Left, distribution of peptides with single (1 lac) or multiple (≥ 2 lac) Klac sites identified via affinity enrichment. Right, distributions of lactyl-NHS–labeled peptides with single (1 lac), double (2 lac), and multiple (> 2 lac) Klac sites at varying lactyl-NHS concentrations in CD8$^+$ T cell lysates, showing lower proportion of single-site labeling at higher concentrations.

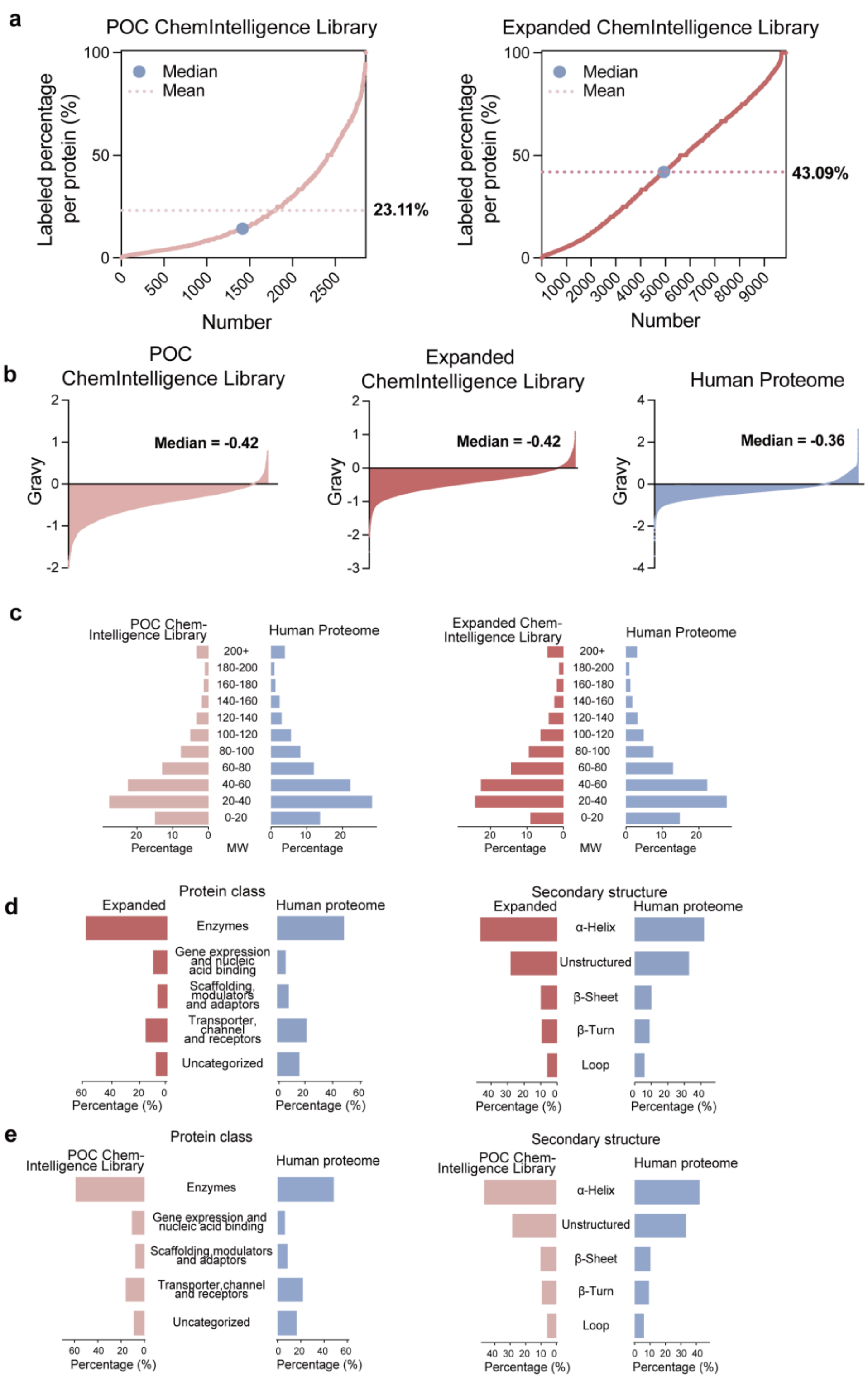


**Supplementary Figure 4. Comparison of biophysical properties between proteins in the Klac ChemIntelligence Library and the human proteome.**

(a) Proportion of lactylated lysines relative to total lysines in each protein across the POC and expanded Klac ChemIntelligence Libraries. The dashed line represents the mean, and the blue dot indicates the median.

(b) Grand average of hydropathy (GRAVY) score distributions in the POC (left), expanded

Klac ChemIntelligence Library (middle) and the human proteome (right).

(c) Molecular weight distributions of proteins in the POC (left) and expanded Klac ChemIntelligence Library (right), compared to the human proteome.

(d) Comparison of protein class (left) and secondary structure distributions (right) between lactylated proteins in the expanded ChemIntelligence Library and the entire human proteome.

(e) Comparison of protein class (left) and secondary structure distributions (right) between lactylated proteins in the POC Klac ChemIntelligence Library and the entire human proteome.

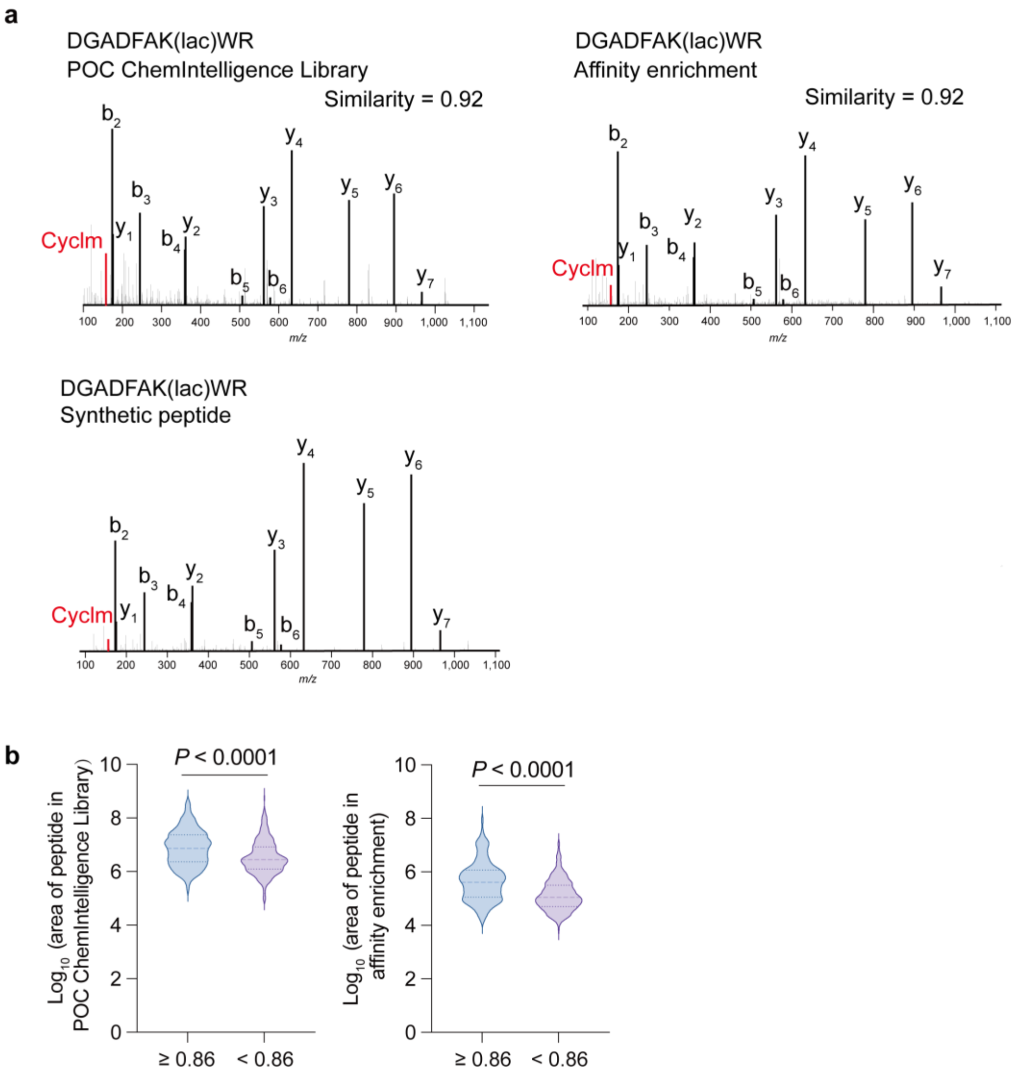


**Supplementary Figure 5. Comparison of MS/MS spectra of Klac peptides from the POC Klac ChemIntelligence Library and affinity enrichment.**

(a) MS/MS spectra of lactyl-NHS–labeled ALDOA-K147lac peptides acquired using three approaches: (1) from the ChemIntelligence Library (top left, Orbitrap Astral), (2) enriched via an affinity antibody-based method (top right, Orbitrap Astral), and (3) a synthetic peptide standard[1] (bottom, Orbitrap Eclipse Tribrid). The cosine similarity between spectra from the ChemIntelligence Library and the antibody-enriched sample is indicated.

(b) Violin plots of peak areas for peptides with spectral cosine similarity $\geq 0.86$ versus $< 0.86$ in the POC Klac ChemIntelligence Library and antibody-enriched samples. Statistical significance was assessed using the Wilcoxon rank-sum test, related to **Fig. 1f**.

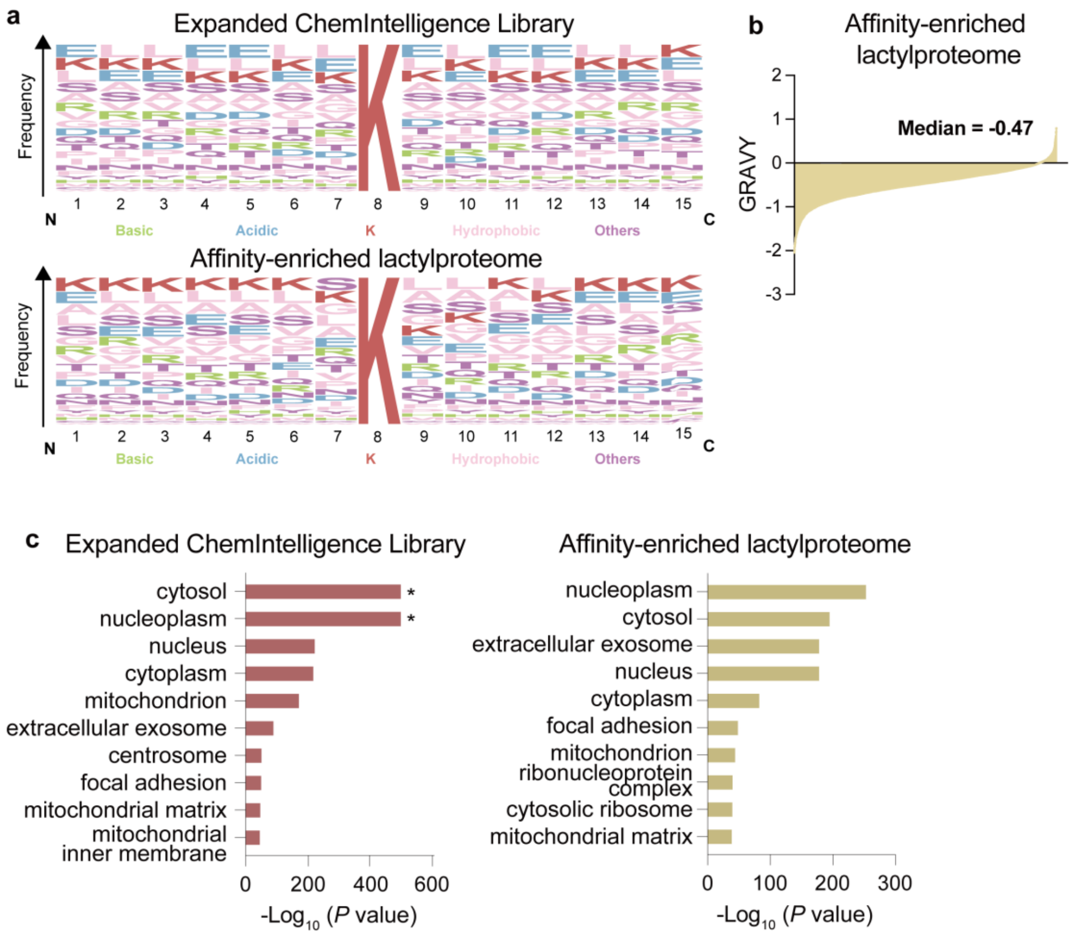


**Supplementary Figure 6. Comparison of lactylated proteins in the expanded Klac ChemIntelligence Library and the affinity-enriched lactylproteome.**

(a) Sequence logos generated using WebLogo showing amino acid frequency surrounding Klac sites in the expanded Klac ChemIntelligence Library (top) and the published affinity-enriched lactylproteome (bottom, related to **Supplementary Table 3**).

(b) GRAVY score distribution of lactylated proteins identified in the published affinity-enriched lactylproteome, related to **Supplementary Fig. 4b**.

(c) GO CC enrichment analysis of lactylated proteins in the expanded Klac ChemIntelligence Library versus the published affinity-enriched lactylproteome (related to **Supplementary Table 3**.). Asterisks indicate *P* values approaching zero, defined as $10^{-500}$ for display purposes.

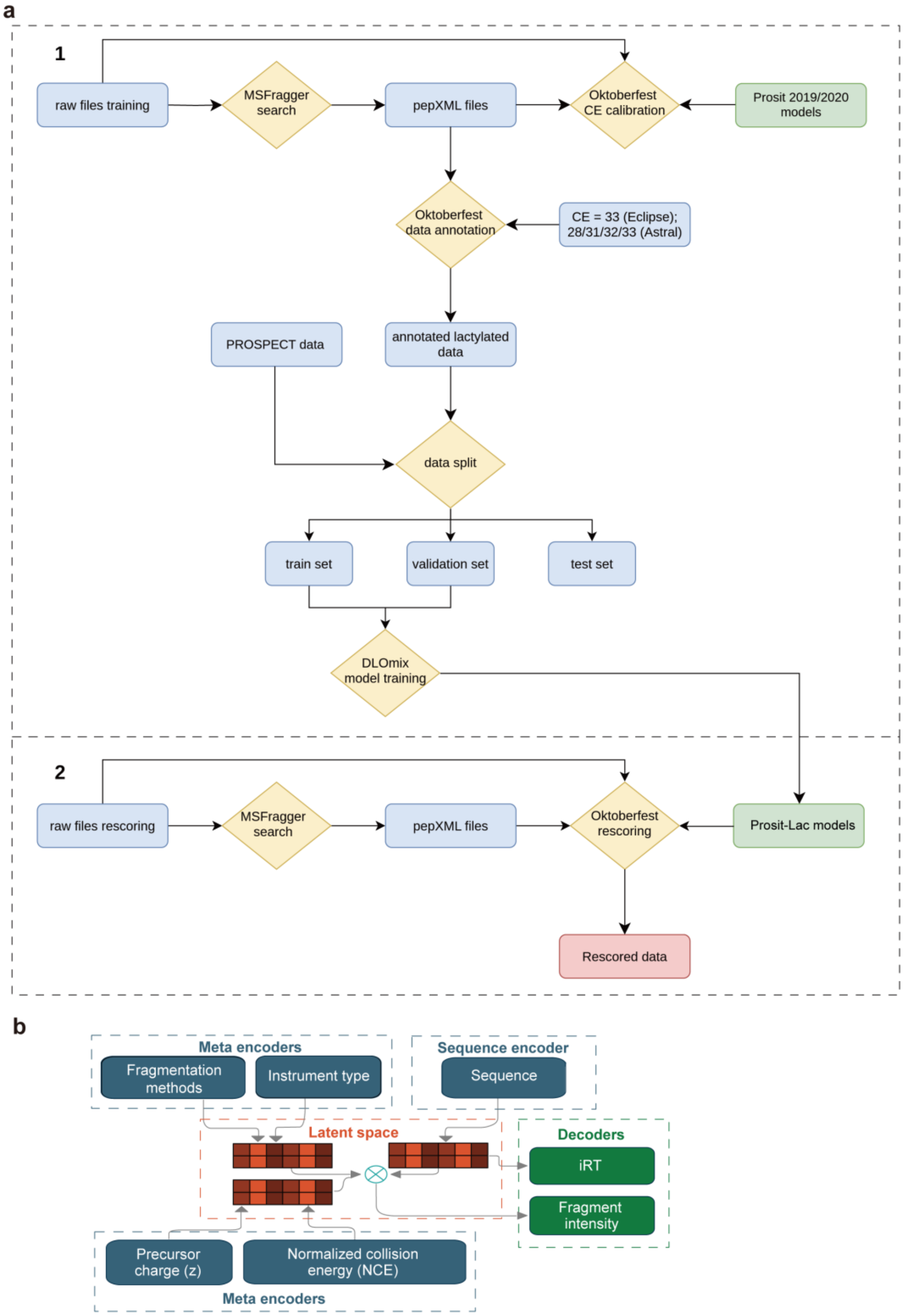
a
1
raw files training
MSFragger search
pepXML files
Oktoberfest CE calibration
Prosit 2019/2020 models
Oktoberfest data annotation
CE = 33 (Eclipse); 28/31/32/33 (Astral)
PROSPECT data
annotated lactylated data
data split
train set
validation set
test set
DLOmix model training
2
raw files rescoring
MSFragger search
pepXML files
Oktoberfest rescoring
Prosit-Lac models
Rescored data
b
Meta encoders
Fragmentation methods
Instrument type
Sequence encoder
Sequence
Latent space
Decoders
iRT
Fragment intensity
Precursor charge (z)
Normalized collision energy (NCE)
Meta encoders

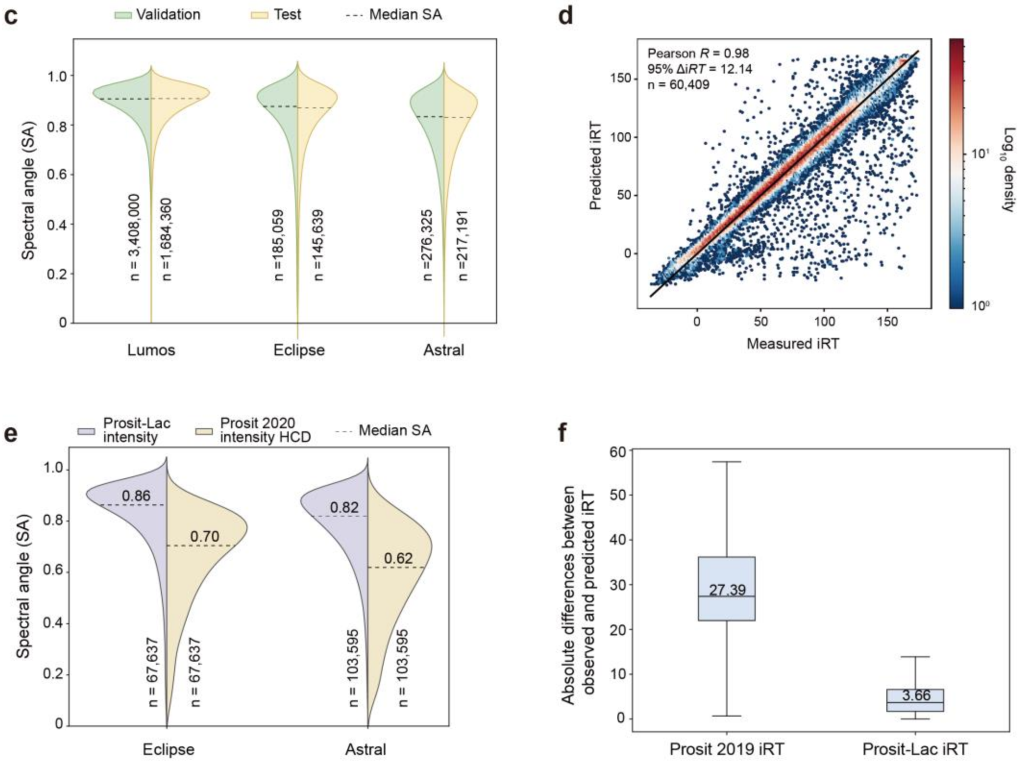


**Supplementary Figure 7. Development and performance evaluation of Prosit-Lac models for Klac peptides.**

(a) Schematic overview of the analysis workflow: (1) Generation of training data and development of the Prosit-Lac models; (2) Rescoring of search engine results using Oktoberfest in combination with the trained Prosit-Lac models.

(b) The Prosit-Lac architecture for fragment ion intensity and iRT prediction.

(c) Violin plots depicting spectral angle distributions of Klac PSMs in the validation set or test set, predicted by the Prosit-Lac model and measured on Orbitrap Tribrid Lumos (Lumos, left), Orbitrap Tribid Eclipse (Eclipse, middle), and Orbitrap Astral (Astral, right) instruments. Green violins represent Klac PSMs in the validation set, while yellow violins represent Klac PSMs in the test set. Dashed lines indicate the median spectral angle within each group.

(d) Scatter density plot of the correlation between Prosit-Lac–predicted and experimentally measured iRTs for the validation set. The color scale indicates point density on a $Log_{10}$ scale.

(e) Violin plots showing the distribution of spectral angles for Klac PSMs predicted by the Prosit-Lac intensity model and by the Prosit 2020 intensity HCD, measured on Eclipse (left) and Astral (right) instruments. Purple violins correspond to predictions from the Prosit-Lac intensity model, while yellow violins correspond to predictions from the Prosit 2020 intensity HCD model. Dashed lines mark the median spectral angle within each group.

(f) Boxplots depicting the absolute differences between observed and predicted iRT values for Klac peptides, comparing the Prosit 2019 iRT (left) and the Prosit-Lac iRT model (right).

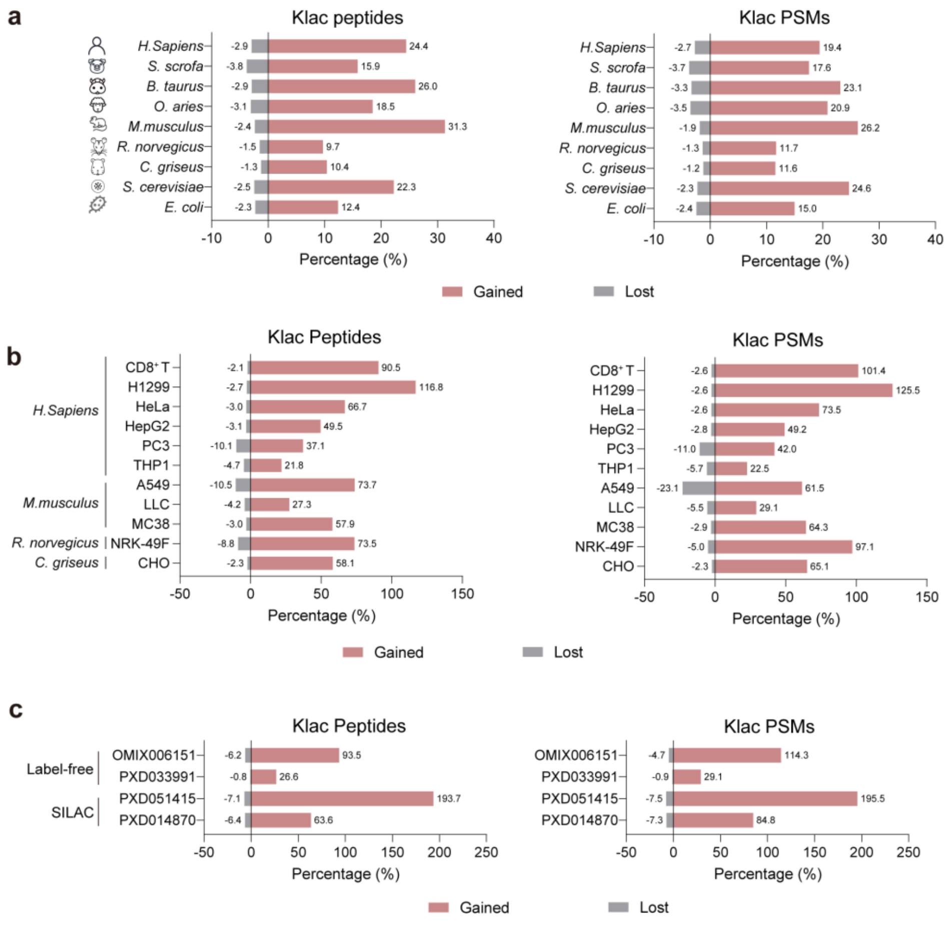


**Supplementary Figure 8. Gains and losses of Klac peptides and PSMs revealed by Oktoberfest rescoring across diverse sample types and species.**

(a) Bar charts showing the percentage of gained (red) and lost (grey) Klac peptides (left) and PSMs (right) after Oktoberfest rescoring of cross-species Klac ChemIntelligence Libraries, benchmarked against the original database search results (related to **Fig. 2c**).

(b) Bar charts showing the percentage of gained (red) and lost (grey) Klac peptides (left) and PSMs (right) after Oktoberfest rescoring of affinity-enriched samples from lactate-treated cell lines of human and animal species, benchmarked against the original database search results (related to **Fig. 2d**).

(c) Bar charts showing the percentage of gained (red) and lost (grey) Klac peptides (left) and PSMs (right) after Oktoberfest rescoring of publicly available affinity-enriched lactylproteome datasets, benchmarked against the original database search results (related to **Fig. 2e**).

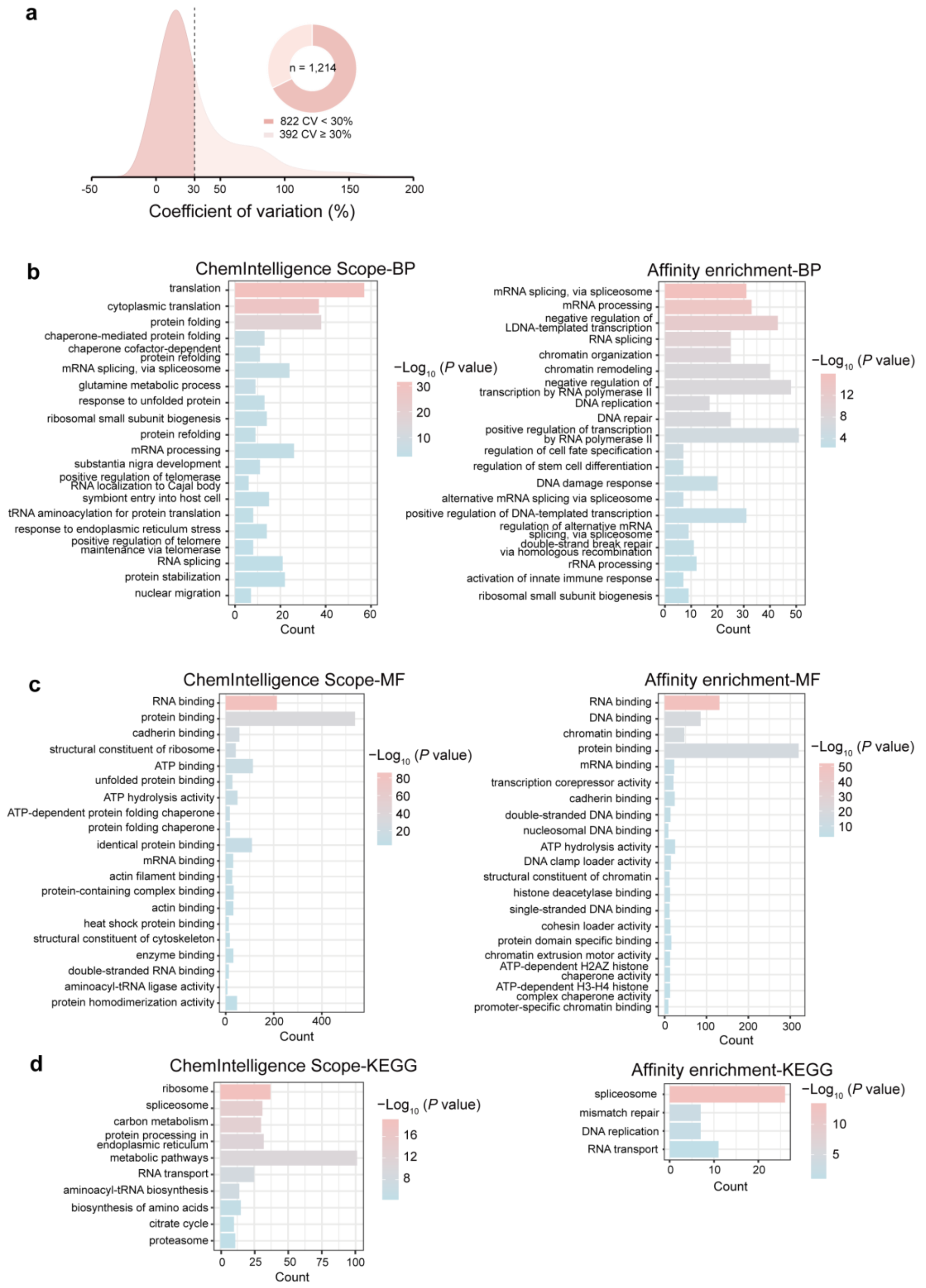


**Supplementary Figure 9. Functional enrichment analyses of Klac proteins uniquely quantified by ChemIntelligence Scope and by affinity enrichment.**

(a) CV distributions of Klac peptides from triplicate ChemIntelligence Scope PRM analyses, with peptides showing CV < 30% (dashed line) considered quantifiable; inset pie chart shows the proportion of peptides across CV categories.

(b-d) Bar charts of GO enrichment analysis showing biological process (BP, b), molecular function (MF, c) and KEGG pathway analysis (d) for proteins uniquely quantified by

ChemIntelligence Scope (left) and by affinity enrichment (right), respectively.

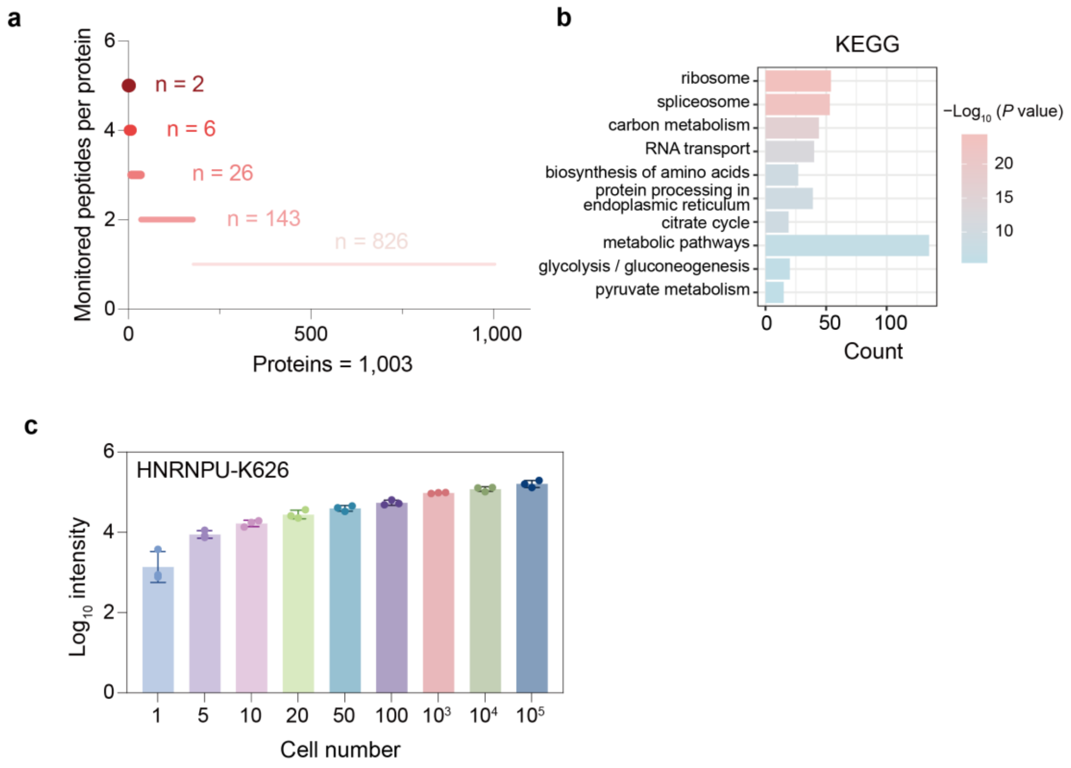


**Supplementary Figure 10. Supporting analyses for single-cell quantification of Klac peptides and proteins.**

(a) Scatter plot showing the distribution of monitored Klac peptides per protein. Each point represents one protein, illustrating that the majority of lactylated proteins contain only a single Klac peptide.

(b) Bar chart of KEGG pathway enrichment analysis for the proteins containing the monitored Klac peptides in (**a**).

(c) Bar chart illustrating the intensity of the lactylate peptide bearing HNRNPU-K626l across samples ranging from 1 to $10^5$ cells (n=3 biological replicates/group), corresponding to **Fig. 4f**.

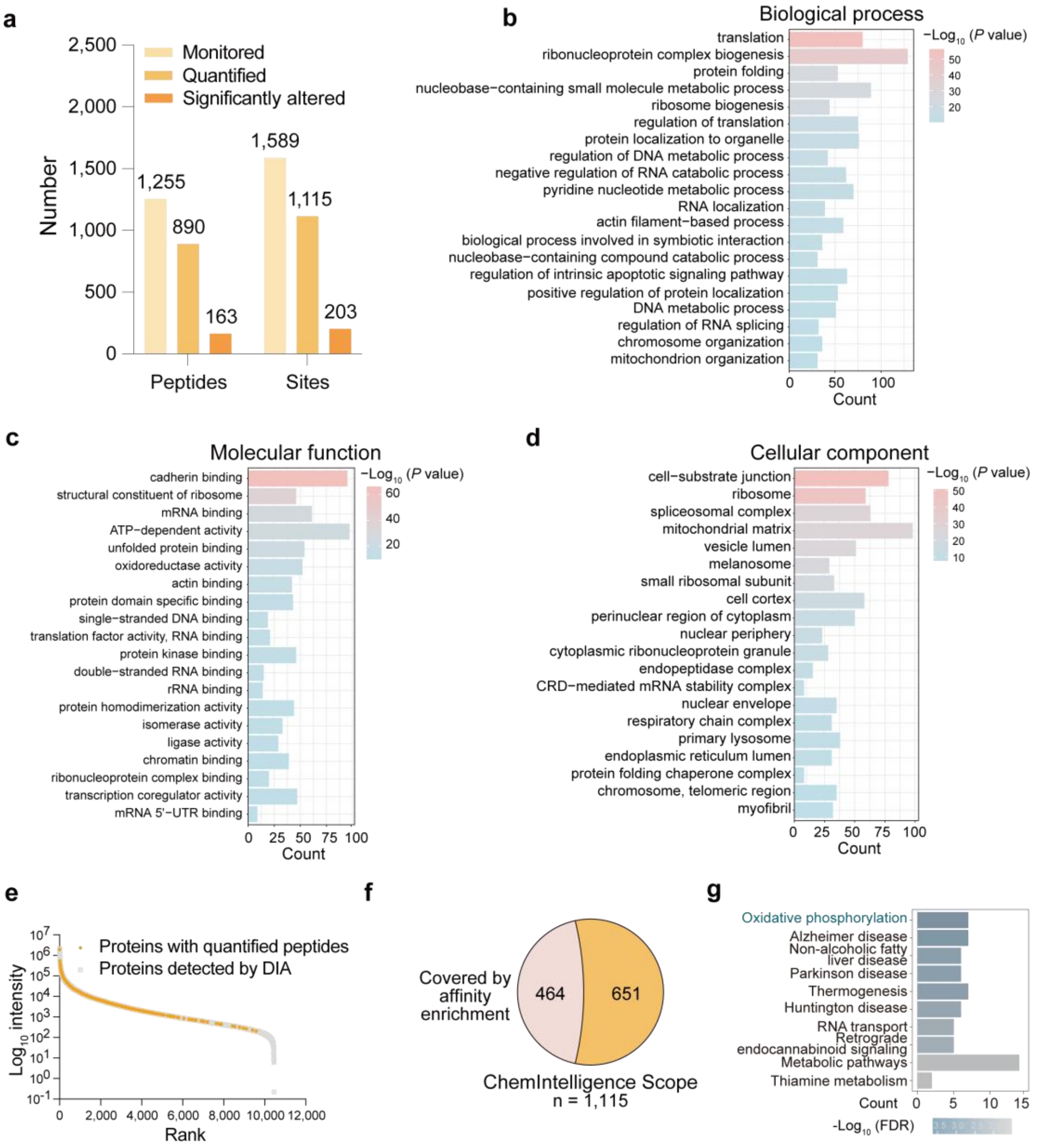
a
Number
2,500
2,000
1,500
1,000
500
0
Monitored
Quantified
Significantly altered
1,255
890
163
1,589
1,115
203
Peptides
Sites
b
Biological process
−Log10 (P value)
50
40
30
20
translation
ribonucleoprotein complex biogenesis
protein folding
nucleobase−containing small molecule metabolic process
ribosome biogenesis
regulation of translation
protein localization to organelle
regulation of DNA metabolic process
negative regulation of RNA catabolic process
pyridine nucleotide metabolic process
RNA localization
actin filament−based process
biological process involved in symbiotic interaction
nucleobase−containing compound catabolic process
regulation of intrinsic apoptotic signaling pathway
positive regulation of protein localization
DNA metabolic process
regulation of RNA splicing
chromosome organization
mitochondrion organization
0
50
100
Count
c
Molecular function
−Log10 (P value)
60
40
20
cadherin binding
structural constituent of ribosome
mRNA binding
ATP−dependent activity
unfolded protein binding
oxidoreductase activity
actin binding
protein domain specific binding
single−stranded DNA binding
translation factor activity, RNA binding
protein kinase binding
double−stranded RNA binding
rRNA binding
protein homodimerization activity
isomerase activity
ligase activity
chromatin binding
ribonucleoprotein complex binding
transcription coregulator activity
mRNA 5'−UTR binding
0
25
50
75
100
Count
d
Cellular component
−Log10 (P value)
50
40
30
20
10
cell−substrate junction
ribosome
spliceosomal complex
mitochondrial matrix
vesicle lumen
melanosome
small ribosomal subunit
cell cortex
perinuclear region of cytoplasm
nuclear periphery
cytoplasmic ribonucleoprotein granule
endopeptidase complex
CRD−mediated mRNA stability complex
nuclear envelope
respiratory chain complex
primary lysosome
endoplasmic reticulum lumen
protein folding chaperone complex
chromosome, telomeric region
myofibril
0
25
50
75
100
Count
e
Log10 intensity
10^7
10^6
10^5
10^4
10^3
10^2
10^1
10^0
10^-1
Proteins with quantified peptides
Proteins detected by DIA
0
2,000
4,000
6,000
8,000
10,000
12,000
Rank
f
Covered by affinity enrichment
464
651
ChemIntelligence Scope
n = 1,115
g
Oxidative phosphorylation
Alzheimer disease
Non-alcoholic fatty liver disease
Parkinson disease
Thermogenesis
Huntington disease
RNA transport
Retrograde endocannabinoid signaling
Metabolic pathways
Thiamine metabolism
Count
0
5
10
15
-Log10 (FDR)

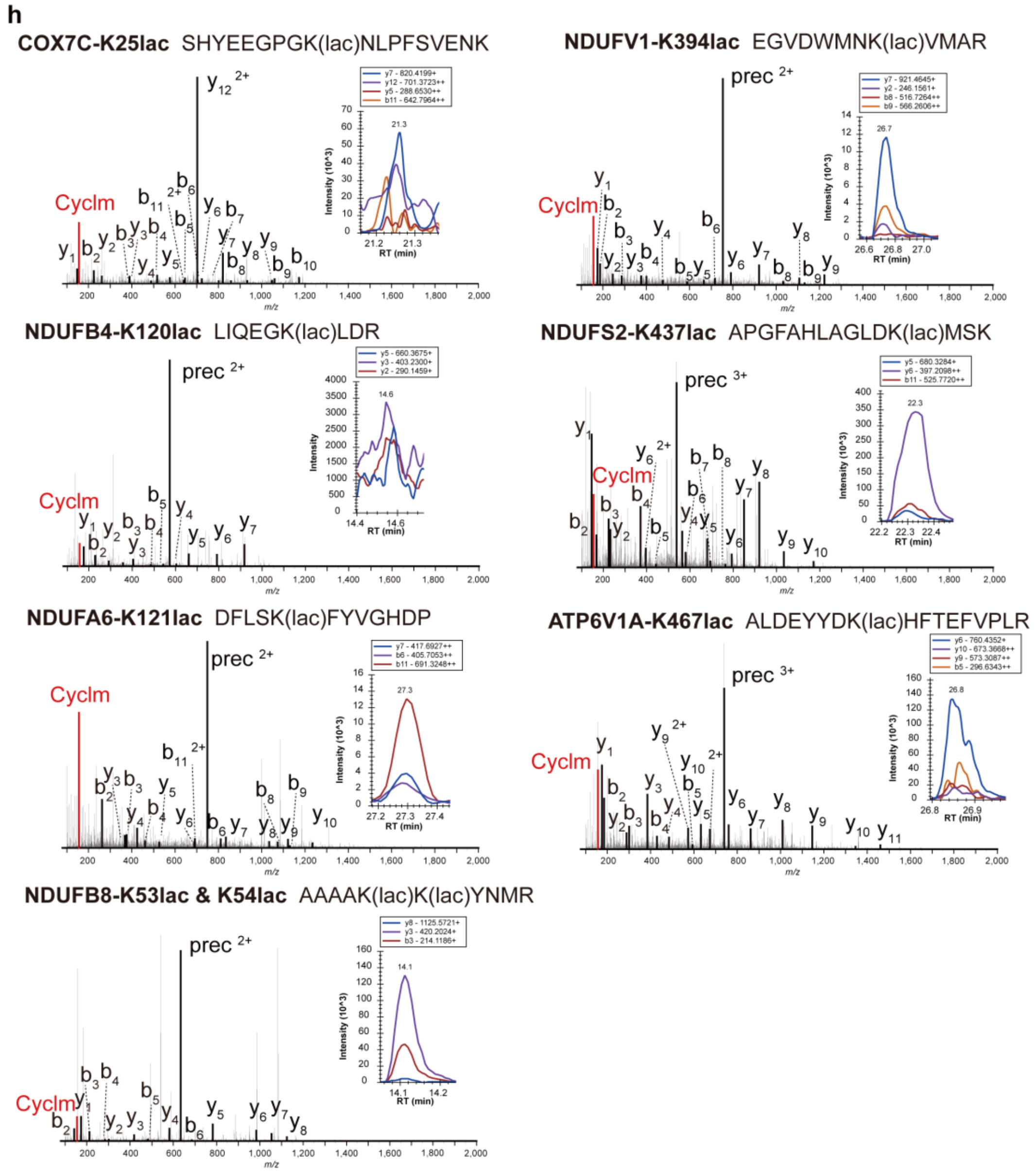


**Supplementary Figure 11. Analysis of quantified Klac peptides, corresponding sites, and proteins in tumor and tumor-adjacent tissue sections of NSCLC patients.**

(a) Bar plot showing the number of Klac peptides/sites monitored, quantified, and significantly altered in three paired NSCLC tumor and adjacent lung sections.

(b-d) Bar charts of GO enrichment analyses showing BP (b), MF (c), and CC (d) categories for proteins with quantified Klac peptides via the ChemIntelligence Scope approach.

(e) Intensity distributions of proteins with quantified peptides in tumor and tumor-adjacent tissue sections. Data represent DIA-based quantitative proteomics of tumor and tumor-adjacent tissue sections from three NSCLC patients, acquired by Orbitrap Astral. Grey, proteins detected by Orbitrap Astral. Orange, proteins with quantified Klac peptides.

(f) Venn diagram showing lactylated sites uniquely quantified in tumor and tumor-adjacent tissue sections using the ChemIntelligence Scope approach, compared to those quantified in affinity-enriched lactylproteome.

(g) KEGG pathway enrichment analysis of proteins with lactylated sites in (**f**) uniquely quantified by ChemIntelligence Scope.

(h) Representative MS/MS spectra and XICs of Klac peptides derived from subunits of mitochondrial respiratory complex I, IV, and V. These peptides were quantifiable by ChemIntelligence Scope in NSCLC tissue sections but absent from previously reported affinity-enriched lactylproteome datasets. MS/MS spectra were retrieved from the POC ChemIntelligence Library, while XICs were extracted from PRM data acquired from paired tissue sections.

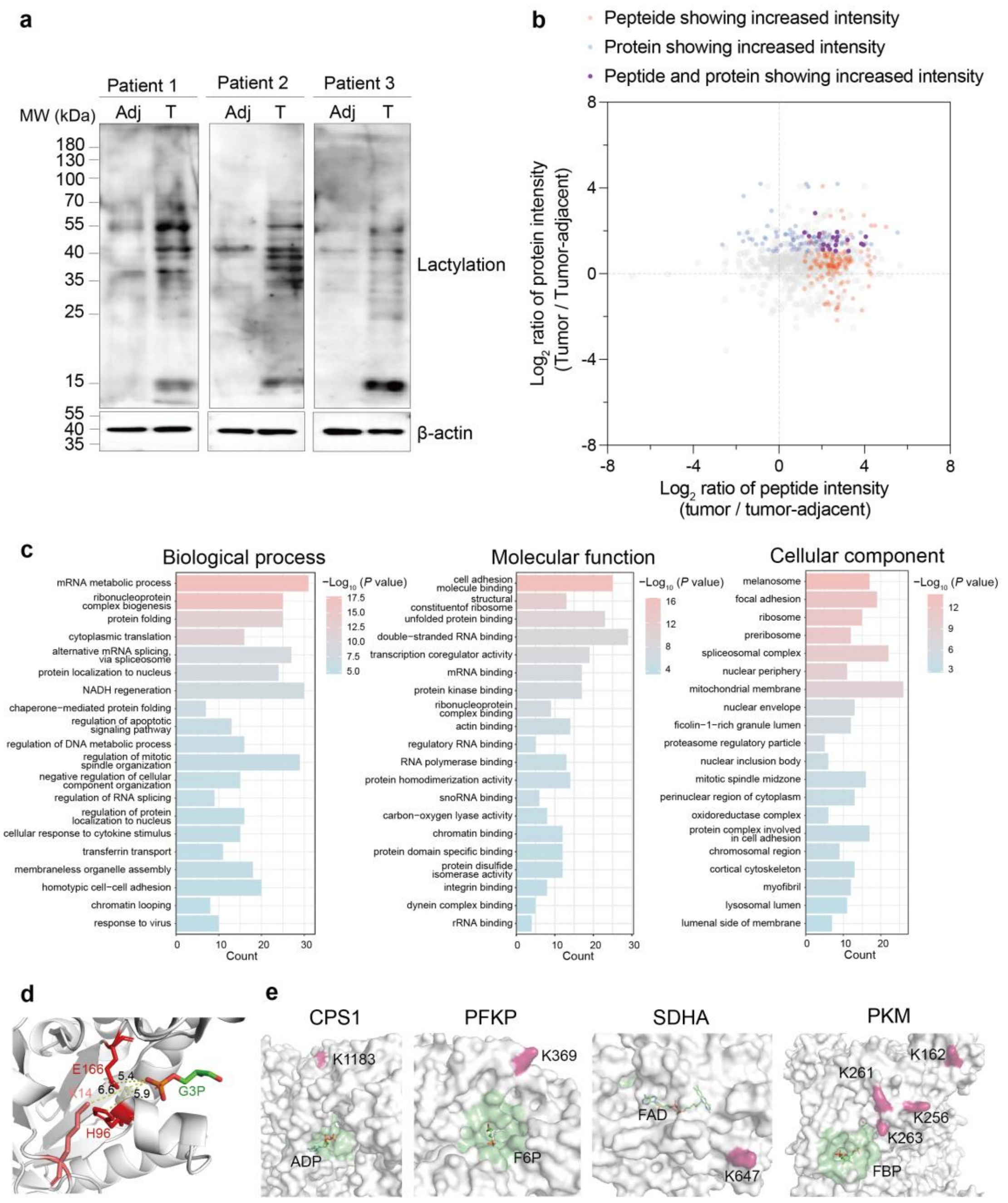


**Supplementary Figure 12. Analysis of elevated Klac peptides, corresponding sites, and proteins in tumor versus tumor-adjacent tissue sections of NSCLC patients.**

(a) Immunoblot analysis showing increased lactylation in tumors compared with adjacent normal tissue sections.

(b) Quadrant analysis of quantified Klac peptide and their protein intensity dynamics between tumor and tumor-adjacent tissue sections. Each point represents one Klac peptide. The x-axis shows the $Log_2$ ratio of peptide intensity (tumor / tumor-adjacent). The y-axis shows the $Log_2$ ratio of protein intensity (tumor / tumor-adjacent). Protein and peptide intensities were acquired using Orbitrap Astral and Stellar MS, respectively. Significantly altered peptides and proteins defined as tumor/tumor-adjacent ratio > 2 or < 0.5, $P$ value < 0.05 by ratio paired $t$-test (n=3 biological replicates/group). Significantly altered peptides and proteins are highlighted in orange and blue, respectively. Klac peptides exhibiting significant changes at both peptide and protein levels are highlighted in purple.

(c) Bar charts of GO enrichment analyses for BP, MF and CC terms of proteins with increased Klac peptides in tumor compared to tumor-adjacent tissue sections.

(d) Crystal structure of TPI1 (PDB: 9FFC) highlighting the lactylated K14 residue located ~6.6 Å from its substrate G3P.

(e) Crystal structures of CPS1 (PDB 5DOU), PFKP (PDB 4XZ2), SDHA (PDB 8GS8) and PKM (PDB 6TTF) showing lactylated sites in pink. Green regions indicate residue within 8 Å of the substrate-binding sites. This spatial arrangement suggests that certain lactylated sites are distal to catalytic centers.

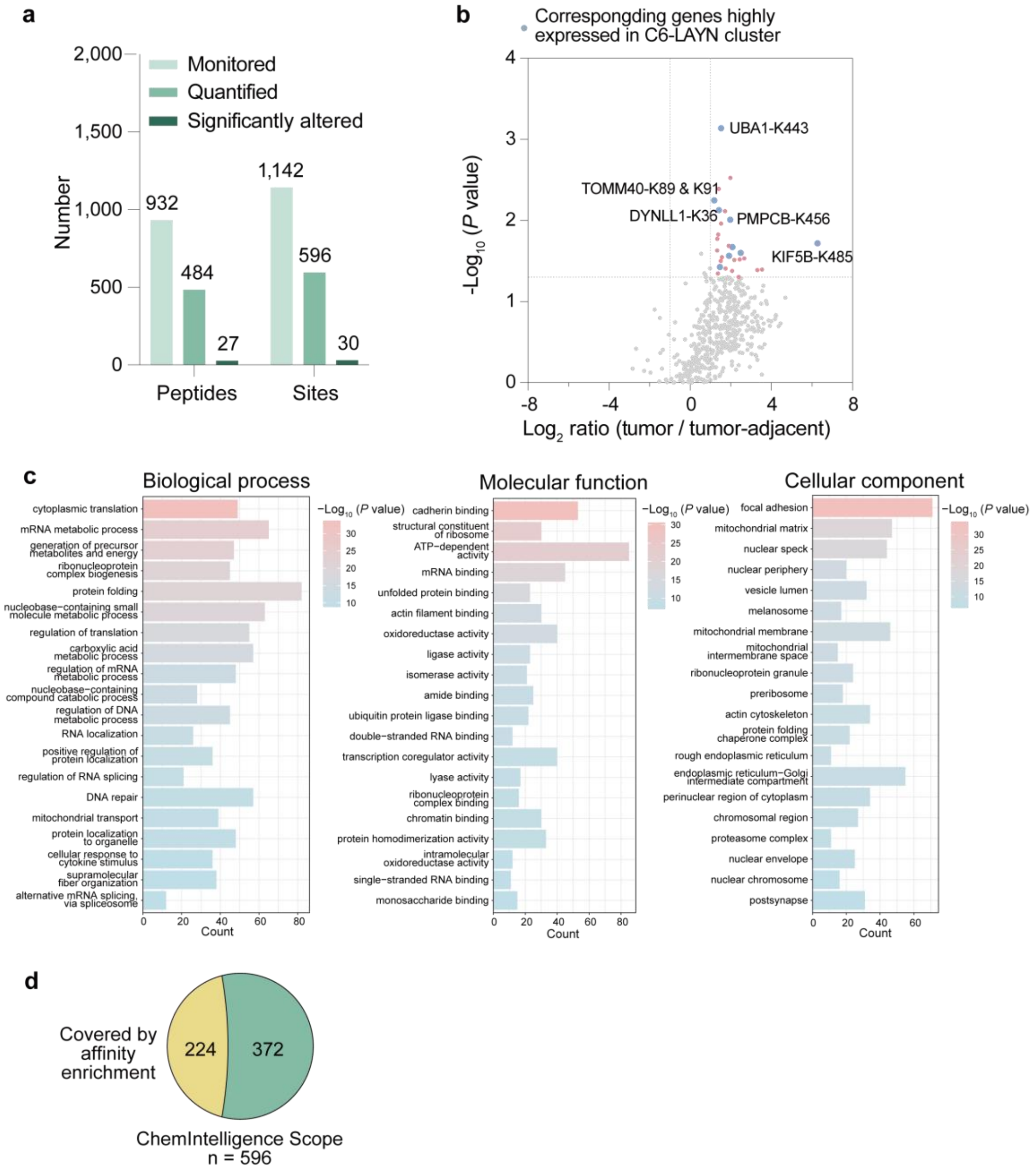


**Supplementary Figure 13. Analysis of quantified Klac peptides, corresponding sites, and proteins in tumor-infiltrating and tumor-adjacent CD8$^+$ T cells from LUAD patients.**

(a) Bar plot showing the number of Klac peptides and sites monitored, quantified, and significantly altered in tumor-infiltrating versus adjacent CD8$^+$ T cells.

(b) Volcano plot comparing Klac peptides between tumor-infiltrating and tumor-adjacent CD8$^+$ T cells. Significantly altered peptides are defined as a tumor/tumor-adjacent ratio > 2 or < 0.5 with *P* value < 0.05 by ratio paired *t*-test (n=3 biological replicates/group) and are highlighted in pink. Upregulated peptides whose matched proteins have high expression in the C6-LAYN cluster are highlighted in blue (related to **Fig. 5h-i**).

(c) Bar charts of GO enrichment analyses for BP, MF and CC terms associated with proteins containing quantified Klac peptides in tumor-infiltrating and tumor-adjacent CD8$^+$ T cells.

(d) Venn diagram showing lactylated sites uniquely quantified in tumor-infiltrating and tumor-adjacent $CD8^+$ T cells using the ChemIntelligence Scope approach, compared to those quantified in the affinity-enriched lactylproteome.

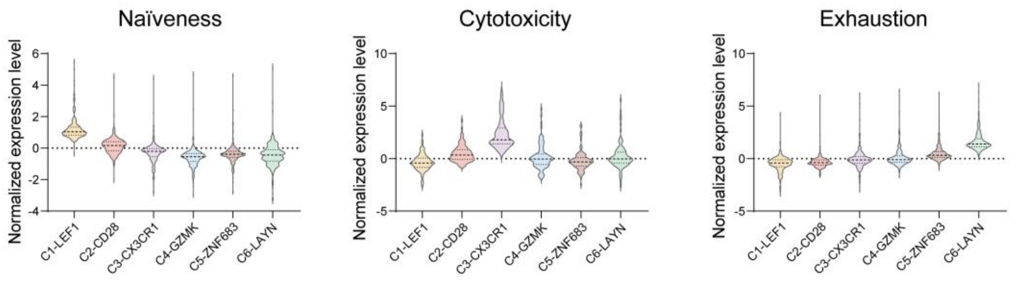


**Supplementary Figure 14. Signature gene expression of CD8⁺ T cells in LUAD patients through re-analysis of single-cell RNA-seq data[2].**

Violin plots showing normalized mean expression of signature genes defining naïve, cytotoxic, and exhausted $CD8^+$ T-cell states across clusters. Data are derived from single-cell RNA-seq of 3,335 T cells isolated from 11 LUAD patients, related to **Fig. 5h.**

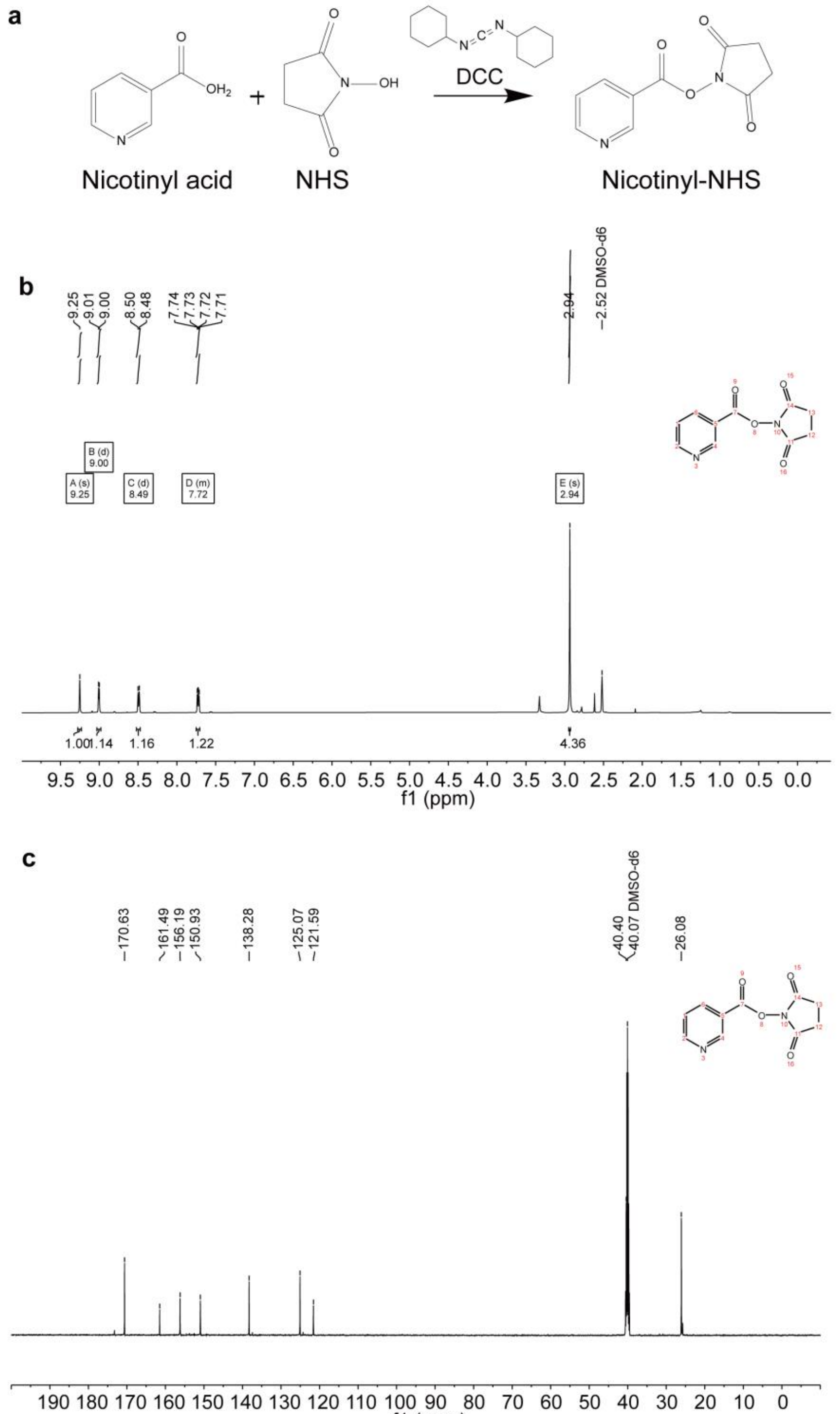


**Supplementary Figure 15. Synthesis and characterization of nicotinyl-NHS.**

(a) Synthetic scheme for generating nicotinyl-NHS ester.

(b) $^{1}H$ NMR spectrum of nicotinyl-NHS in DMSO-$d_6$.

(c) $^{13}C$ NMR spectrum of nicotinyl-NHS in DMSO-$d_6$.

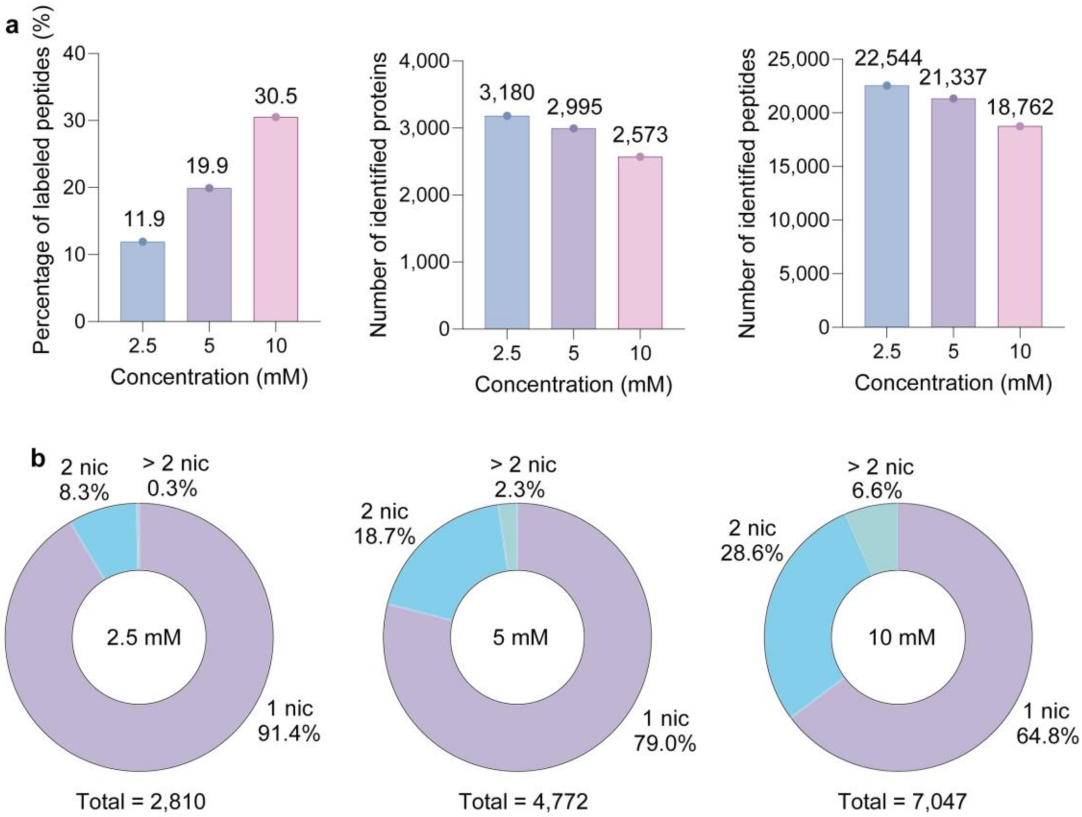


**Supplementary Figure 16. Reactivity of nicotinyl-NHS on cell lysates.**

(a) Percentage of nicotinyl-NHS–labeled peptides relative to total identified peptides (left), total identified proteins (middle) and peptides (right) in HeLa cell lysates at indicated nicotinyl-NHS concentrations.

(b) Distributions of nicotinyl-NHS–labeled peptides with single (1 lac), double (2 lac), or multiple (> 2 lac) Knic sites in HeLa cell lysates at varying nicotinyl-NHS concentrations, showing a reduced proportion of single-site labeling at higher concentrations.

### Supplementary Tables

**Table 1.** Klac peptides identified in the proof-of-concept Klac ChemIntelligence Library.

**Table 2.** Klac peptides identified in the expanded Klac ChemIntelligence Library.

**Table 3.** Summary of Klac sites from the affinity-enriched lactylproteome reported in previous studies.

**Table 4.** Summary of 21 PRM assays performed on Stellar, targeting Klac peptides from the proof-of-concept Klac ChemIntelligence Library.

**Table 5.** Summary of Klac peptides quantified by ChemIntelligence Scope in control and lactate–treated HeLa cells.

**Table 6.** Summary of 5 PRM assays targeting Klac peptides newly identified by Oktoberfest rescoring across three technical replicates.

**Table 7.** Summary of 19 PRM assays targeting Klac peptides from the ultra-low-input Klac ChemIntelligence Library.

**Table 8.** Summary of Klac peptides monitored by ChemIntelligence Scope in lactate–treated HeLa cells across a range from single-cell to $10^5$-cell inputs in three biological replicates.

**Table 9.** Clinical characteristics of NSCLC patients included in this study.

**Table 10.** Klac peptides quantified by population-based pre-screening in H1299 cells.

**Table 11.** Summary of Klac peptides quantified by ChemIntelligence Scope in tumor and adjacent tissue sections from NSCLC patient samples.

**Table 12.** Summary of Klac peptides quantified by population-based pre-screening in $CD8^+$ T cells.

**Table 13.** Summary of Klac peptides quantified by ChemIntelligence Scope in tumor-infiltrating and tumor-adjacent $CD8^+$ T cells from LUAD patient samples.

**Table 14.** Summary of Knic peptides identified in the Knic ChemIntelligence Library.

**Table 15.** Summary of 24 PRM assays targeting Knic peptides from the Knic ChemIntelligence Library.

**Table 16.** Summary of Knic peptides quantified by ChemIntelligence Scope-Knic in control and nicotinate–treated HeLa cells.
